# CDK9 and PP2A regulate RNA polymerase II transcription termination and coupled RNA maturation

**DOI:** 10.1101/2021.06.21.449289

**Authors:** Michael Tellier, Justyna Zaborowska, Jonathan Neve, Takayuki Nojima, Svenja Hester, Marjorie Fournier, Andre Furger, Shona Murphy

## Abstract

CDK9 is a kinase critical for the productive transcription of protein-coding genes by RNA polymerase II (pol II). As part of P-TEFb, CDK9 phosphorylates the carboxyl-terminal domain (CTD) of pol II and elongation factors, which allows pol II to elongate past the early elongation checkpoint (EEC) encountered soon after initiation. We show that, in addition to halting pol II at the EEC, loss of CDK9 activity causes premature termination of transcription across the last exon, loss of polyadenylation factors from chromatin, and loss of polyadenylation of nascent transcripts. Inhibition of the phosphatase PP2A abrogates the premature termination and loss of polyadenylation caused by CDK9 inhibition, indicating that this kinase/phosphatase pair regulates transcription elongation and RNA processing at the end of protein-coding genes. We also confirm the splicing factor SF3B1 as a target of CDK9 and show that SF3B1 in complex with polyadenylation factors is lost from chromatin after CDK9 inhibition. These results emphasize the important roles that CDK9 plays in coupling transcription elongation and termination to RNA maturation downstream of the EEC.

## Introduction

Transcription of human protein-coding genes by RNA polymerase (pol) II is a complex process comprising initiation, elongation, and termination. In addition, mRNA capping, splicing, and cleavage and polyadenylation, which are required to produce a mature mRNA, are largely co-transcriptional (Tellier, Maudlin et al., 2020a). The dynamic phosphorylation and dephosphorylation of proteins that control transcription and pre-mRNA processing, including pol II itself, are fundamental to the regulation of gene expression. Phosphorylation of pol II mostly occurs on the carboxyl-terminal domain (CTD) of its largest subunit, RBP1, which comprises 52 repeats of the heptapeptide sequence Tyr1,Ser2,Pro3,Thr4,Ser5,Pro6,Ser7 (YSPTSPS) (Zaborowska, Egloff et al., 2016). During transcription, tyrosine, serine and threonine residues are reversibly and dynamically phosphorylated whilst the two prolines are subject to cis-trans isomerisation. The pattern of heptapeptide phosphorylation and proline isomerisation during the transcription process creates distinct CTD profiles throughout the transcription cycle to coordinate the recruitment of transcription elongation and pre-mRNA processing factors in space and time (Buratowski, 2009, Corden, 2013, Hsin & Manley, 2012, Zaborowska et al., 2016). For example, CTD Ser5 phosphorylation helps to recruit capping proteins at the 5’ end of genes and Ser2 phosphorylation helps to recruit polyadenylation and termination factors at the 3’ end of genes.

In human cells, phosphorylation of the pol II CTD heptapeptide is mainly carried out by cyclin-dependent kinases (CDKs), including CDK7, CDK9, and CDK12 (Zaborowska et al., 2016). CDK9, together with Cyclin T1, forms the Positive Transcription Elongation Factor b complex (P-TEFb), which phosphorylates Ser2, Thr4, and Ser5 of the CTD heptapeptide. In addition, P-TEFb phosphorylates several proteins involved in regulation of transcriptional elongation, including the negative elongation factor subunit E (NELFE), the SPT5 subunit of DSIF, and SPT6 (Peterlin & Price, 2006, Vos, Farnung et al., 2018a, Vos, Farnung et al., 2018b, Yamada, Yamaguchi et al., 2006). Soon after initiation of transcription of protein-coding genes, pol II stalls at an early elongation checkpoint (EEC) due to the recruitment of NELF and DSIF. Phosphorylation of these complexes by P-TEFb results in release of NELF and turns DSIF into a positive elongation factor, allowing productive elongation to proceed (Jonkers, Kwak et al., 2014, Laitem, Zaborowska et al., 2015, Vos et al., 2018a, Vos et al., 2018b). Accordingly, inhibition of CDK9 by small molecule inhibitors causes pol II to pause at the EEC genome-wide (Jonkers et al., 2014, Laitem et al., 2015, Vos et al., 2018a, Vos et al., 2018b). Using short-term treatment of cells with CDK9 inhibitors, we have shown that inhibition of CDK9 also disrupts transcription at the 3’end of protein-coding genes, causing premature termination of pol II close to the poly(A) site (Laitem et al., 2015). This premature termination is associated with the loss of several pol II-associated factors, including CDK9 itself, SPT5, SSU72, and Cstf64 from the 3’ end of protein-coding genes (Laitem et al., 2015). This led us to hypothesize the presence of a poly(A)-associated checkpoint (PAAC), where CDK9 is required to overcome the pause and transcribe past the poly(A) site for cleavage and polyadenylation to occur (Laitem et al., 2015, Tellier, Ferrer-Vicens et al., 2016).

Termination of transcription downstream of an active polyadenylation site requires the action of the exonuclease Xrn2, which degrades the uncapped RNA left associated with pol II after cleavage of the nascent pre-mRNA at the poly(A) site (Proudfoot, 2016). This is thought to destabilize pol II to allow termination of transcription. Interestingly, phosphorylation of Xrn2 by CDK9 enhances its activity (Sanso, Levin et al., 2016).

As the functions of transcriptional kinases are becoming clearer, thanks in part to the development of cell lines with analog-sensitive kinases (Bishop, Ubersax et al., 2000, Tellier, Zaborowska et al., 2020b), the role of phosphatases in this process are beginning to emerge. Several CDK9 targets, including SPT5 and Xrn2, are dephosphorylated by the multifunctional protein phosphatases (PP)1, PP2A, and PP4 (Huang, Jee et al., 2020, Parua, Booth et al., 2018, Parua, Kalan et al., 2020, Vervoort, Welsh et al., 2021), which also have roles in splicing (Mermoud, Cohen et al., 1992, Shi, Reddy et al., 2006). In addition, PP1 was recently shown to regulate poly(A)-site-dependent transcription termination by dephosphorylation of SPT5 (Cortazar, Sheridan et al., 2019, Eaton, Francis et al., 2020). Knockdown of PP1 or its associated subunit, PNUTS, or inhibition of PP1 with the small molecule inhibitor Tautomycetin, leads to hyperphosphorylation of SPT5 and the elongation rate of pol II no longer decreases downstream of the poly(A). This leads to a transcription termination defect as the exonuclease Xrn2 fails to “catch up” with the elongating pol II (Cortazar et al., 2019, Eaton et al., 2020). PP2A interacts with Integrator, a protein complex that cleaves RNA and regulates the amount of paused pol II at the EEC (Huang et al., 2020, Vervoort et al., 2021, Zheng, Qi et al., 2020). PP2A is also thought to dephosphorylate the pol II CTD and other factors involved in the early stages of transcription (Huang et al., 2020, Vervoort et al., 2021, Zheng et al., 2020).

There is also evidence that termination of transcription of intron-containing protein-coding genes in human cells is coupled to terminal exon definition, which requires the coordinated recognition of the 3’SS and the poly(A) site (Cooke, Hans et al., 1999, Tellier et al., 2020a). For example, mutation of the 3’SS causes a transcription termination defect due to failure to recognise the poly(A) site (Dye & Proudfoot, 1999). Interactions between splicing factors, such as SF3B1 or U2AF65, and CPA factors, such as CPSF100 or poly(A) polymerase, PAPOLA, have been demonstrated (Gunderson, Beyer et al., 1994, Kyburz, Friedlein et al., 2006, Niwa, Rose et al., 1990, Tellier et al., 2020a, Vagner, Vagner et al., 2000). In addition, subunit CDC73 of the PAF complex, which plays a role in elongation, interacts with the CPSF73 CPA endonuclease (Rozenblatt-Rosen, Nagaike et al., 2009). Thus, inhibition of CDK9 may trigger premature termination by disrupting the balance of kinase and phosphatase activities required for enabling protein-protein/-DNA/-RNA interactions.

To elucidate the function of CDK9 in the PAAC, we generated, using CRISPR/Cas9, a HEK293 cell line where the endogenous genes express analog-sensitive CDK9 (CDK9as). Inhibition of CDK9as or small molecule CDK9 inhibitors promote premature termination of pol II, impair CPA factor recruitment to chromatin, and cause loss of polyadenylation of newly-made pre-mRNA. Interestingly, bioinformatic analysis indicates that the defect in transcription caused by CDK9 inhibition starts across the last exon, rather than at the poly(A) site, implicating disruption of definition of the last exon in premature termination.

Using phosphoproteomics, we have identified numerous targets of CDK9 including, as expected, SPT5 and NELF, in addition to the U2 snRNP splicing factor component SF3B1, with SF3B1 T142P confirmed as a PP1 target. Inhibition of CDK9 does not affect interaction between CPA factors and the SF3B complex but rather promotes the loss of interaction between pol II and a complex containing SF3B and CPA factors, suggesting that CDK9 plays a role in the recruitment of these factors to the elongation complex at the end of genes. PP1 inhibition promotes a transcription termination defect, as previously shown (Eaton et al., 2020) but does not reverse the effect of CDK9 inhibition on transcription and pol II CTD phosphorylation. However, inhibition of PP2A reverses the effect of CDK9 inhibition on premature termination, indicating that this phosphatase has a previously unsuspected role at the 3’end of protein-coding genes. Inhibition of PP2A also restores recruitment of poly(A) factors and the polyadenylation of newly-made pre-mRNA disrupted by CDK9 inhibition. PP2A inhibition alone causes an increase in pol II signal across the last exon, a greater recruitment of SF3B1 to the pol II complex, and a higher production of polyadenylated mRNA indicating that PP2A is a negative regulator of mRNA CPA.

Taken together our results highlight an additional kinase/phosphatase switch at the end of protein-coding genes regulating the coordination of pre-mRNA splicing, cleavage/polyadenylation, and termination of transcription.

## Results

### CDK9 inhibition abrogates polyadenylation

We have previously shown that CDK9 inhibition leads to loss of pol II association downstream of the poly(A) site of human protein-coding genes (Laitem et al., 2015). This premature termination of transcription could be accompanied by loss of polyadenylation and failure to make a mature mRNA. Alternatively, polyadenylation/termination may be more efficient and fully mature mRNA produced. We have already shown that levels of the polyadenylation factor CstF64 at the 3’ end of genes is reduced after CDK9 inhibition, suggesting that polyadenylation is compromised. In order to follow the production of newly-polyadenylated mRNA, we have analysed production of transcripts from TNFα-inducible genes. HeLa cells were treated with DMSO or TNFα for 30 minutes followed by treatment with DMSO or 5,6- dichlorobenzimidazone-1-β-D-ribofuranoside (DRB) for 30 minutes and 3’READS was then carried out on the purified nuclear mRNAs to measure the production of newly polyadenylated mRNA (Figure 1A) (Neve, Burger et al., 2016). There are 307 genes where mRNA production is induced more than two-fold by TNFα in both repeats. Pol II ChIP-qPCR on one of these genes, *LDLR*, was carried out to analyse induction of transcription by TNFα and the effect of DRB treatment on pol II at the 3’end (Appendix Figure S1A). Polyadenylation of the 307 TNFα-induced mRNAs in the nucleus is significantly decreased, both for all of the genes included and for genes longer than 40 kb, where pol II is still elongating at the 3’end (Figure 1B and Appendix Figure S1A). qRT-PCR of the nuclear polyadenylated mRNAs encoded by selected TNFα-induced genes confirmed that DRB causes a marked reduction in polyadenylation (Appendix Figure S1B).

**Figure 1.**
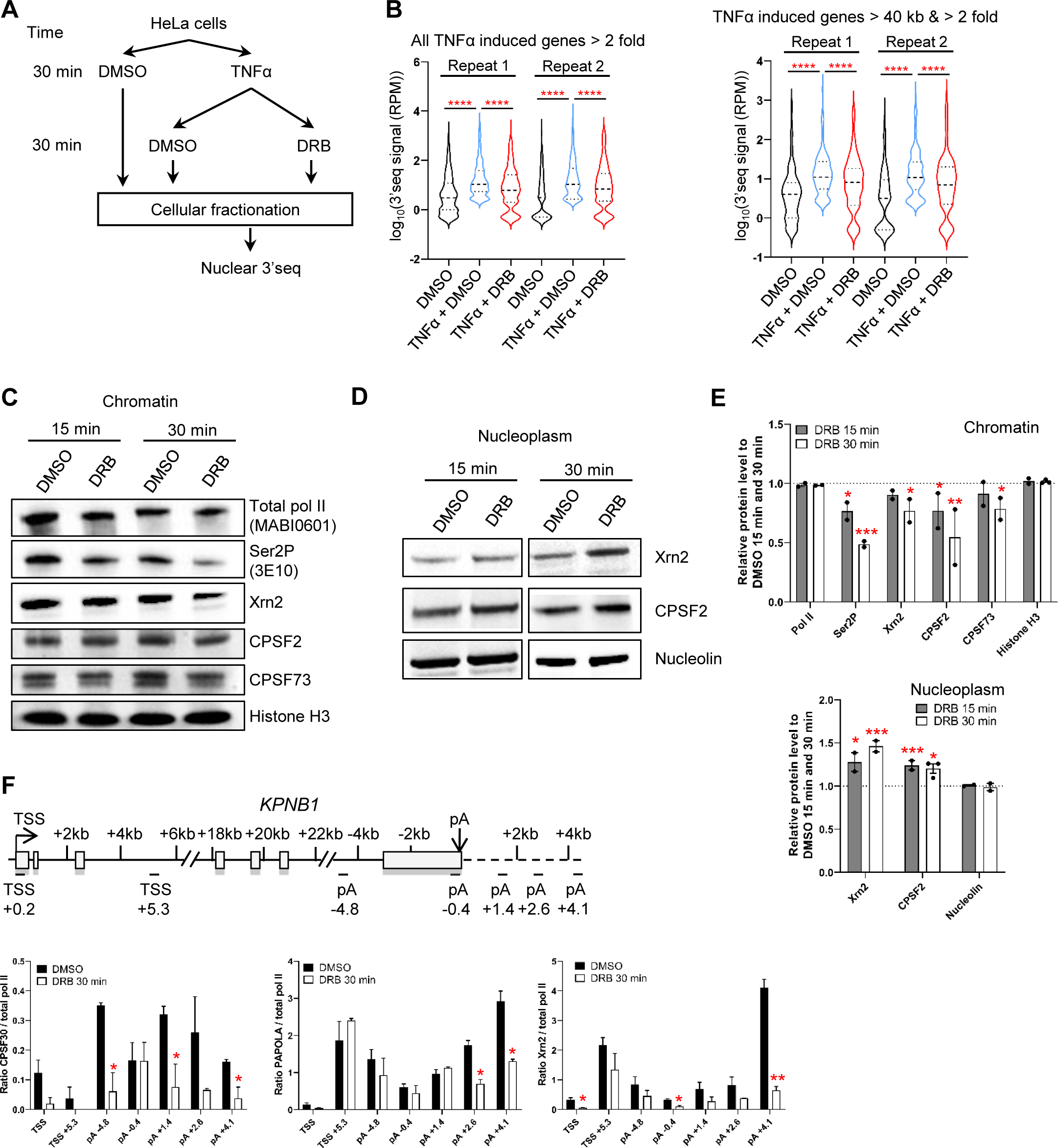
CDK9 inhibition abrogates polyadenylation. **A.** Schematic of the nuclear 3’READS experiment. **B.** Violin plots of the 3’READS signal on all the TNFα induced genes (n=307) or the set of TNFα induced genes longer than 40 kb (n=111). **** p < 0.0001. Statistical test: Friedman test with Dunn’s multiple test correction. **C.** Western blot of total pol II, Ser2P, Xrn2, CPSF2, CPSF73, and histone H3 as a loading control, on the chromatin fraction. **D.** Western blot of Xrn2, CPSF2, and Nucleolin as a loading control, on the nucleoplasm fraction (the CPSF73 antibody does not provide reliable results on the nucleoplasm fraction). **E.** Quantification of the western blots shown in **C** and **D**. n=2 biological replicates, mean ± SEM, p-value: * p < 0.05, ** p < 0.01, *** p < 0.001. Statistical test: two-tailed unpaired t test. **F.** Ratio of the different CPA and termination factors to total pol II. n=3 biological replicates, mean ± SEM, p-value: * p < 0.05, ** p < 0.01. Statistical test: two-tailed unpaired t test.

We also performed western blotting to assess the level of the polyadenylation/termination factors Xrn2, CPSF2, CPSF73, total pol II, and CTD Ser2P in chromatin and nucleoplasm after treatment of HeLa cells for 15 or 30 minutes with DRB (Figure 1C-E and Appendix Figure S1C).

As expected, Ser2P is decreased on the chromatin following CDK9 inhibition. There is also a loss of Xrn2, CPSF2 and CPSF73 from the chromatin fraction, with a stronger loss after 30 than 15 minutes. In contrast, the level of these factors in the nucleoplasm fraction increases, indicating that these factors dissociate from chromatin after CDK9 inhibition. Importantly, there is no reduction of these factors in whole cell extract, indicating that the loss from chromatin is not due to active degradation (Appendix Figures S1D and S1E). Levels of the poly(A) polymerase (PAPOLA), Xrn2 and CPSF30, measured by ChIP-qPCR on our model gene, *KPNB1*, which is approximately 35 kb long, are also reduced after 30 minutes treatment with DRB, whether ratioed to pol II levels or not (Figure 1F and Appendix Figure S1F).

This data indicates that CDK9 inhibition causes both premature termination of pol II and failure to recruit polyadenylation factors, which causes the production of polyadenylated mRNA to be aborted.

### CDK9 inhibition causes an elongation defect starting at the last exon of protein-coding genes

To better understand the kinetics of premature termination of pol II caused by CDK9 inhibition, mNET-seq was carried out with a total pol II antibody after treatment of HeLa cells with DRB for 5, 10, 15, or 30 minutes (Figure 2A-C and Figure EV1A and EV1B). The results are similar to those we previously obtained using GRO-seq (Laitem et al., 2015), and metagene profile analysis indicates that DRB treatment of cells causes an increase in pol II pausing close to the TSS, a loss of pol II entering productive elongation, and premature termination of pol II close to the poly(A) site is detected on genes longer than 40 kb where pol II has not yet “run off” (Figures 2B-D, and Figure EV1A and EV1B). Pol II ChIP-qPCR on *KPNB1* after treatment of cells with DRB concentrations between 12.5 to 100 µM gives the same result (Figure EV1C). Interestingly, these data also show that CDK9 inhibition affects pol II at the EEC more rapidly (easily detectable after 5 minutes inhibition) than at the 3’end of genes (only detectable after 10 minutes inhibition).

**Figure 2.**
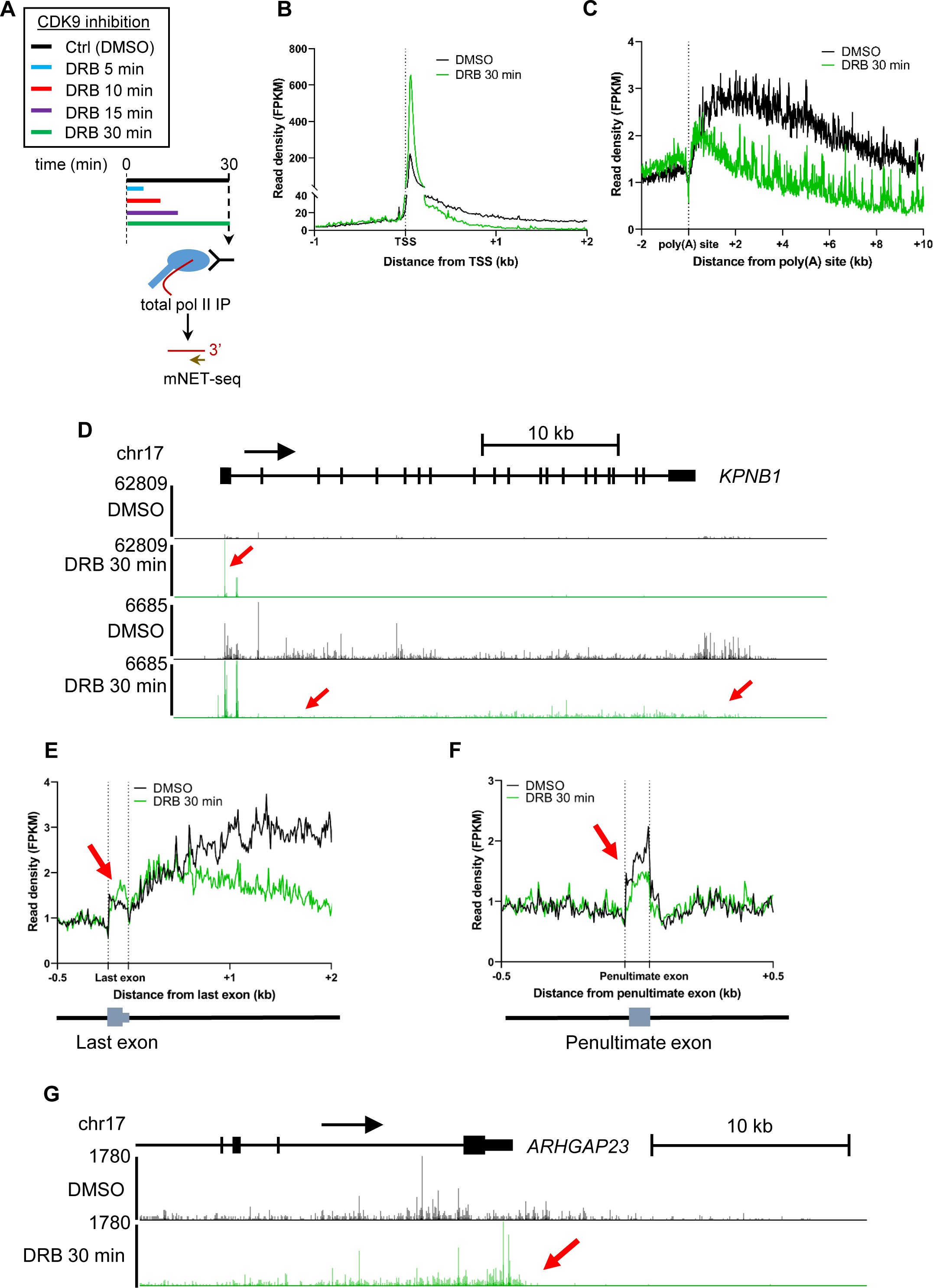
CDK9 inhibition causes an elongation defect starting at the last exon of protein-coding genes. **A.** Schematic of the total pol II mNET-seq experiments. **B.** Metagene profile of total pol II after 30 minutes treatment with DMSO (black) or DRB (green) around the TSS of expressed protein-coding genes (n=6,965). **C.** Metagene profile of total pol II after 30 minutes treatment with DMSO (black) or DRB (green) around the used poly(A) site of expressed protein-coding genes longer than 40 kb (n=2,816). **D.** Screenshot of the genome browser mNET-seq tracks across the protein-coding gene *KPNB1*. Red arrows indicate increased pol II pausing, loss of pol II entering productive elongation, and premature termination. **E.** Metagene profile of total pol II after 30 minutes treatment with DMSO (black) or DRB (green) around the used last exon of expressed protein-coding genes (n=2,526). **F.** Metagene profile of total pol II after 30 minutes treatment with DMSO (black) or DRB (green) around the penultimate exon of expressed protein-coding genes (n=2,426). **G.** Screenshot of the genome browser mNET-seq tracks at the 3’end of protein-coding gene *ARHGAP23*. The red arrow indicates increased pol II signal across the last exon.

To focus on the effect of CDK9 inhibition on pol II behaviour at the 3’ end of genes, a metagene analysis was carried out with the pol II signal scaled across the last exon (Figure 2E). Interestingly, the metaprofile indicates that CDK9 inhibition causes an increase in pol II signal across the last exon followed by loss of pol II signal downstream of the poly(A) site. Importantly, the increase in the pol II signal after CDK9 inhibition is specific to the last exon as the pol II signal over the penultimate exons or other internal exons is either unchanged or decreased (Figure 2F and Figure EV1D). This is particularly apparent on, for example, *ARHGAP23* where pol II levels increase specifically across the last exon (Figure 2G).

These findings indicate that the defect in transcription at the 3’ end of genes caused by CDK9 inhibition starts upstream of the poly(A) site and occurs more slowly than loss of transcription at the EEC.

### Inhibition of analog-sensitive (as) CDK9 has the same effect on elongation as small molecule CDK9 inhibitors

Small molecule kinase inhibitors may target more than one kinase *in vivo* (Bensaude, 2011). To confirm that the results we observed with DRB are due solely to inhibition of CDK9, we used CRISPR/Cas9 to change the gatekeeper phenylalanine to an alanine in all endogenous copies of CDK9 HEK293 cells to make a CDK9 analog-sensitive (as) cell line (Figure EV2A), which has a similar growth rate to the wild type cells (Figure EV2B). Treatment of wild-type HEK293 or CDK9as cells with 7.5, 10, or 15 µM of the ATP-analogue 1-NA-PP1 (NA), which should specifically inhibit CDK9as (Zhang, Lopez et al., 2013), affects the growth rate of the CDK9as cells with little effect on the wild-type HEK293 cells (Figure EV2C). Pol II ChIP-qPCR of *KPNB1* in CDK9as cells after treatment of cells with 7.5, 10, or 15 µM NA for 15 minutes indicates that at all NA concentrations, pol II pausing and pol II entry into productive elongation is affected (Supplementary Figure 3-1D). Readily detectable premature termination of pol II close to the poly(A) site is caused by 15 µM NA (Figure EV2D). Importantly, 15 µM NA has no effect on pol II in wild-type HEK293 (Figure EV2E). Introduction of the CDK9as mutation or treatment with 15 µM NA also do not affect the level of CDK9 or Cyclin T1 in whole cell extract (Figures EV2F and EV2G). NA treatment of wild-type HEK293 cells also has no effect on polyadenylation (Figure EV2H).

The effect of CDK9 inhibition on pol II CTD phosphorylation was analysed by western blotting of chromatin fractions (Figures 3A and B). As expected, CDK9as inhibition causes loss of Ser2P and Ser5P with two different phosphoantibodies to each CTD mark (ab5095 and ab5121 from Abcam, 13499S and 13523S from Cell Signaling) while NA treatment has no effect on CTD phosphorylation in wild-type HEK293 cells. ChIP-qPCR on *KPNB1* also indicates that CDK9as inhibition causes loss of Ser2P and Ser5P, whether ratioed to pol II or not (Figure 3C and Appendix Figure S2), indicating that *in vivo* CDK9 activity is necessary for efficient Ser2 and Ser5 phosphorylation.

**Figure 3.**
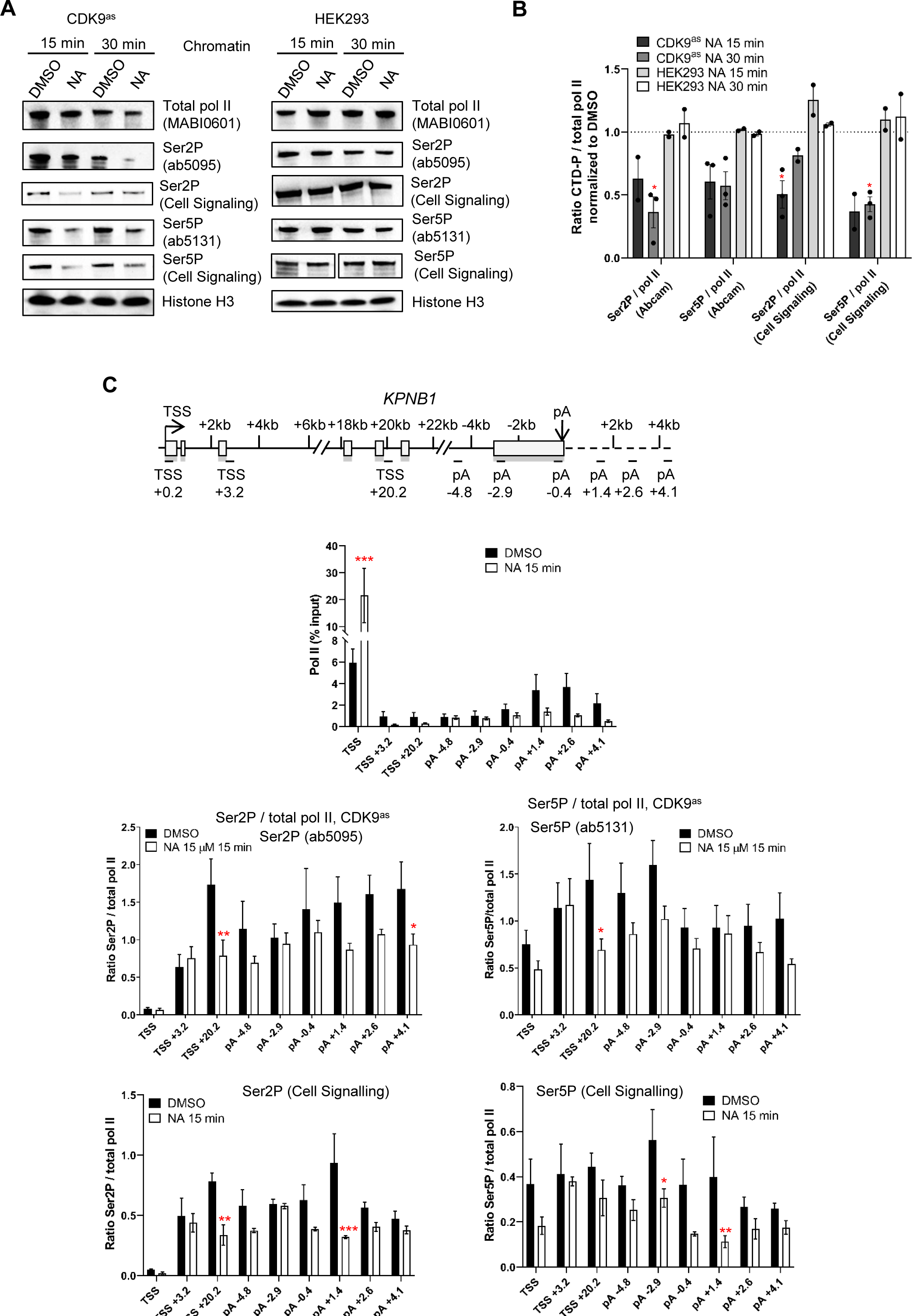
Inhibition of analog-sensitive (as) CDK9 has the same effect on elongation as small molecule CDK9 inhibitors. **A.** Western blot of total pol II, Ser2P (Abcam or Cell Signaling), Ser5P (Abcam or Cell Signaling), and histone H3 as a loading control, on the chromatin fraction of CDK9as or wild-type HEK293 cells treated with DMSO or 15 µM 1-NA-PP1 for 15 or 30 minutes. **B.** Quantification of the western blots shown in A. n=3 biological replicates, mean ± SEM, p-value: * p < 0.05. Statistical test: two-tailed unpaired t test. **C.** ChIP-qPCR of total pol II and of Ser2P (Abcam or Cell Signaling), and Ser5P (Abcam or Cell Signaling) ratioed to total pol II on *KPNB1*. n=3 biological replicates, mean ± SEM, p-value: * p < 0.05, ** p < 0.01, *** p < 0.001. Statistical test: two-tailed unpaired t test.

These findings reinforce the conclusion that the effect of DRB on pol II is due to CDK9 inhibition.

### CDK9 phosphorylates several transcription and splicing factors *in vivo*

Phosphorylation of several non-CTD CDK9 targets plays critical roles in gene expression (Decker, Forne et al., 2019, Sanso et al., 2016, Zaborowska et al., 2016). To identify the *in vivo* targets of CDK9, we performed biological duplicates of SILAC phosphoproteomics in HeLa cells treated with DMSO or DRB for 30 minutes (Figure 4A and Figure EV3A, Appendix Table 3). We found 100 phosphosites across 74 proteins decreased more than 1.5 fold and with a p-value < 0.1. Amongst these phosphosites, 34 have a SP or a TP motif and are located in 25 different proteins (Figure EV3B). In line with previous similar studies, most CDK9 targets are factors involved in transcription, RNA processing, and RNA biology. Importantly, we identified several known targets of CDK9, including MED1 (T1440), MEPCE (T213, S217, T291), NELFA (S225), SPT5 (S666, S671), and SPT6 (T1523) (Decker et al., 2019, Sanso et al., 2016, Vos et al., 2018a, Vos et al., 2018b). We also identified several residues of the splicing factors SF3B1 (T142, T227, T436) and CDC5L (T377, T396, T430, T442) as targets of CDK9. However, there is only limited overlap with previously published CDK9 *in vitro* or *in vivo* phosphoproteomics (Figure EV3C) (Decker et al., 2019, Sanso et al., 2016).

**Figure 4.**
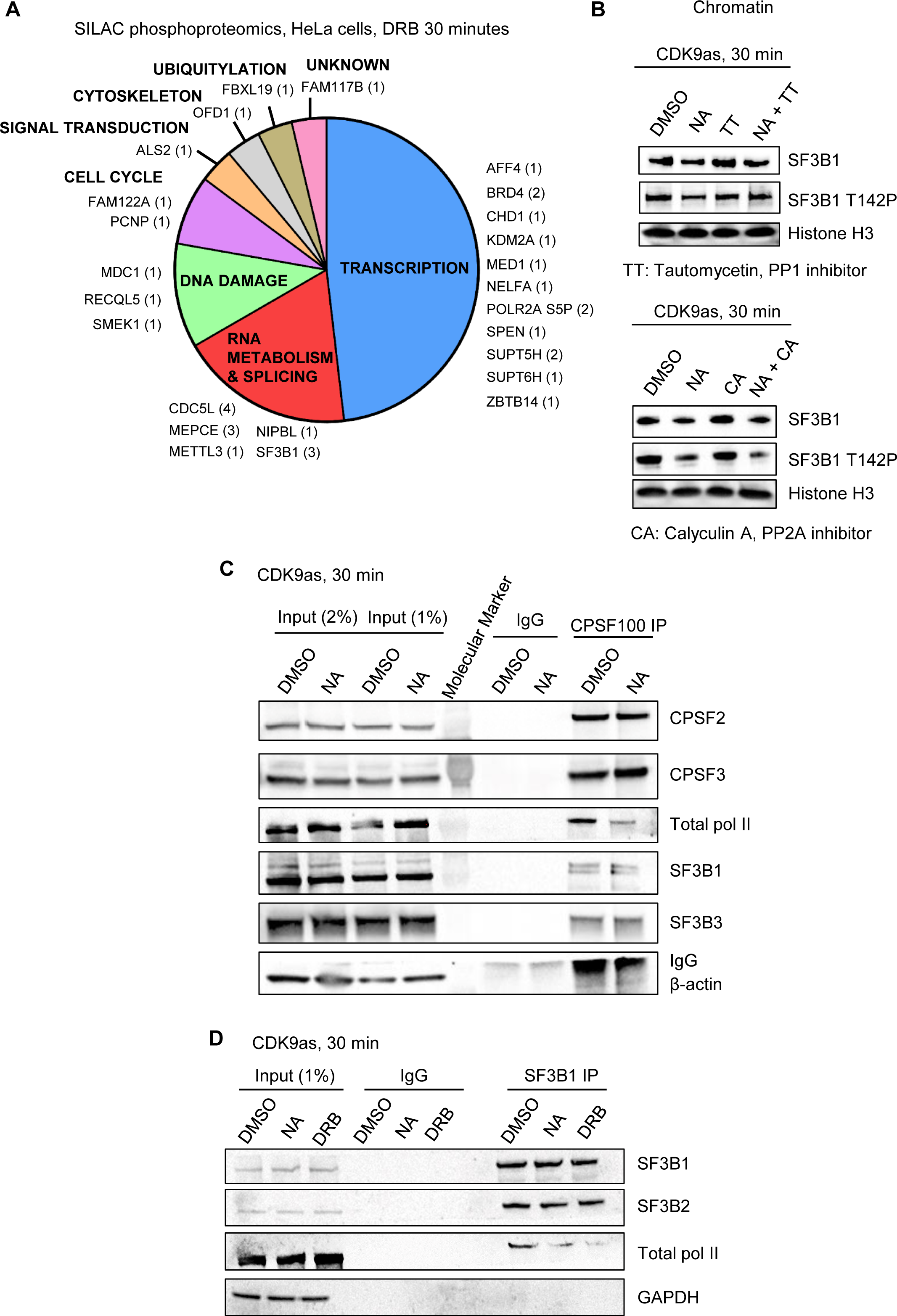
CDK9 phosphorylates several transcription and splicing factors *in vivo*. **A.** Distribution of the proteins found to be phosphorylated by CDK9 in SILAC phosphoproteomics in HeLa cells treated or not with 100 µM DRB for 30 minutes (fold change > 1.5 in both biological duplicates, p-value < 0.1, number of phosphopeptides decreased shown in brackets). **B.** Western blot of total SF3B1, SF3B1 T142P, and histone H3 as a loading control, on the chromatin fraction of CDK9as cells treated for 30 minutes with DMSO, NA, tautomycetin (TT), calyculin A (CA), NA+TT, or NA+CA. **C.** Co-immunoprecipitation of CPSF2 in the CDK9as cell treated for 30 minutes with DMSO or NA followed by western blot with total pol II, CPSF2, CPSF3, SF3B1, SF3B3 and β-actin (negative control) antibodies. **D.** Co-immunoprecipitation of SF3B1 CPSF2 in the CDK9as cell treated for 30 minutes with DMSO or NA followed by western blot with total pol II, SF3B1, SF3B2, and GAPDH (negative control) antibodies.

We have generated a new phosphoantibody for SF3B1 T142P, as SF3B1 is known to have functions in splicing and definition of the last exon (Kyburz et al., 2006, Tellier et al., 2020a). Kinase-phosphatase switches have recently been shown to regulate transcription at the 3’ end of gene and CDK9 and PP1 activities play a major role in controlling SPT5 T806 phosphorylation and pol II elongation close to poly(A) sites (Cortazar et al., 2019, Parua et al., 2018, Parua et al., 2020). In addition, PP2A is known to dephosphorylate SPT5 S666P (Huang et al., 2020, Parua et al., 2018, Parua et al., 2020). Accordingly, we analysed the effect on transcription of low concentrations of three small molecule inhibitors, 25 nM Tautomycetin (TT), which inhibits PP1, and 2.5 nM Calyculin A (CA) or 2.5 nM LB-100, which inhibits PP2A. H3S10P and Myc S62P, which are known targets of PP1 and PP2A, respectively, were used to control for specific inhibition (Appendix Figure S3A). We observed that at the concentrations used, TT preferentially inhibits PP1, while CA and LB-100 mainly inhibit PP2A.

Western blotting of the chromatin fraction of cells treated or not with CDK9 inhibitors confirms that CDK9 phosphorylates SF3B1 T142P while PP1, but not PP2A, dephosphorylates this phosphoresidue (Figure 4B and Figure EV3D).

These findings expand the repertoire of transcription and RNA processing factors phosphorylated by CDK9 and emphasize the important role that reversible protein phosphorylation plays in the regulation of transcription and co-transcriptional processes at the end of genes.

### CDK9 inhibition promotes the loss of interaction of SF3B1 and CPA factors with pol II

The SF3B complex has been shown to interact with mRNA CPA factors to promote mRNA cleavage and polyadenylation (Kyburz et al., 2006). Accordingly, we investigated whether the loss of SF3B1 phosphorylation following CDK9 inhibition could result in a loss of interaction with CPA factors, which could in turn explain the loss of mRNA polyadenylation after CDK9 inhibition. SF3B1 immunoprecipitation from HeLa cells did not indicate that CPA factors strongly interact with SF3B1 (Appendix Table 4). However, as it is possible that only a small proportion of SF3B1 interacts with CPA factors, we performed CPSF2 immunoprecipitation from the CDK9as cell line with or without NA treatment (Figure 4D). Interaction between CPSF2 and CPSF3, SF3B1, and SF3B3 was detected. Interestingly, the interaction between CPSF and SF3B proteins was not affected by NA treatment. However, the interaction between CPSF2 and pol II was markedly decreased by CDK9 inhibition. There is also decreased interaction between SF3B1 and pol II following CDK9 inhibition as shown by immunoprecipitation of SF3B1 from the CDK9as cell line with or without NA or DRB treatment (Figure 4E). Importantly, the interaction between SF3B1 and SF3B2 is not affected by NA or DRB treatment. These experiments suggest that the loss of SF3B1 from chromatin (Figure 4B and Figures EV3E) is due to loss of interaction with pol II.

Thus, CDK9 inhibition decreases SF3B1 phosphorylation and its interaction with total pol II without affecting SF3B1 interaction with other SF3B proteins and CPA factors.

### PP2A counteracts CDK9 activity at the 3’ end of genes

As PP1 and PP2A reverse CDK9-mediated phosphorylation of several residues of SPT5 and SF3B1 (Figure 4B) (Cortazar et al., 2019, Parua et al., 2018, Parua et al., 2020) the effect of PP1 and PP2A inhibition on pol II CTD Ser2 and Ser5 phosphorylation in the absence or presence of CDK9as inhibition was investigated (Appendix Figures S3B and S3C). As expected, NA causes loss of Ser2P and Ser5P while inhibition of PP1 with TT or PP2A with CA results in an increase of Ser2P and Ser5P, in line with the known roles of PP1 and PP2A as both CTD Ser2 and Ser5 phosphatases (Vervoort et al., 2021, Washington, Ammosova et al., 2002, Zheng et al., 2020). TT in addition to NA does not reverse the decrease in Ser2P and Ser5P levels seen in CDK9as cells, whereas CA in addition to NA causes an increase in Ser2P and Ser5P levels compared to NA alone, indicating that, for CTD phosphorylation, PP1 inhibition cannot overcome CDK9 inhibition whereas PP2A inhibition can.

The effect of inhibition of these two phosphatases on transcription of the *KPNB1* gene was therefore investigated by pol II ChIP-qPCR (Figure 5A and Appendix Figure S4A). Inhibition of PP1 leads to an increase in pol II pausing at the TSS coupled to a termination defect downstream of the poly(A) site (primer TerDef), as previously reported (Eaton et al., 2020). Inhibition of PP2A by CA or LB-100 also results in an increase in pol II pausing at the TSS with apparent termination of pol II closer to the poly(A) site (CA, primer TerDef). Inhibition of CDK9 and PP1 at the same time has the same effect as CDK9 inhibition alone (Figure 5B). However, inhibition of PP2A and CDK9 at the same time mitigates the effect of CDK9 inhibition at the 3’end of the gene (see primer pA +1.4 for both DRB + CA and DRB + LB-100). Inhibition of CDK9 for 15 minutes followed by treatment with CA for another 15 minutes, or vice-versa, gives a similar outcome, although there is an increased level of pol II in the gene body when DRB is followed by CA (Appendix Figure S4B). A similar approach with TT results in the same outcome as DRB treatment alone (Appendix Figure S4C). Treatment of cells with DRB, CA, and TT together has the same effect as DRB and CA (Figure 5C).

**Figure 5.**
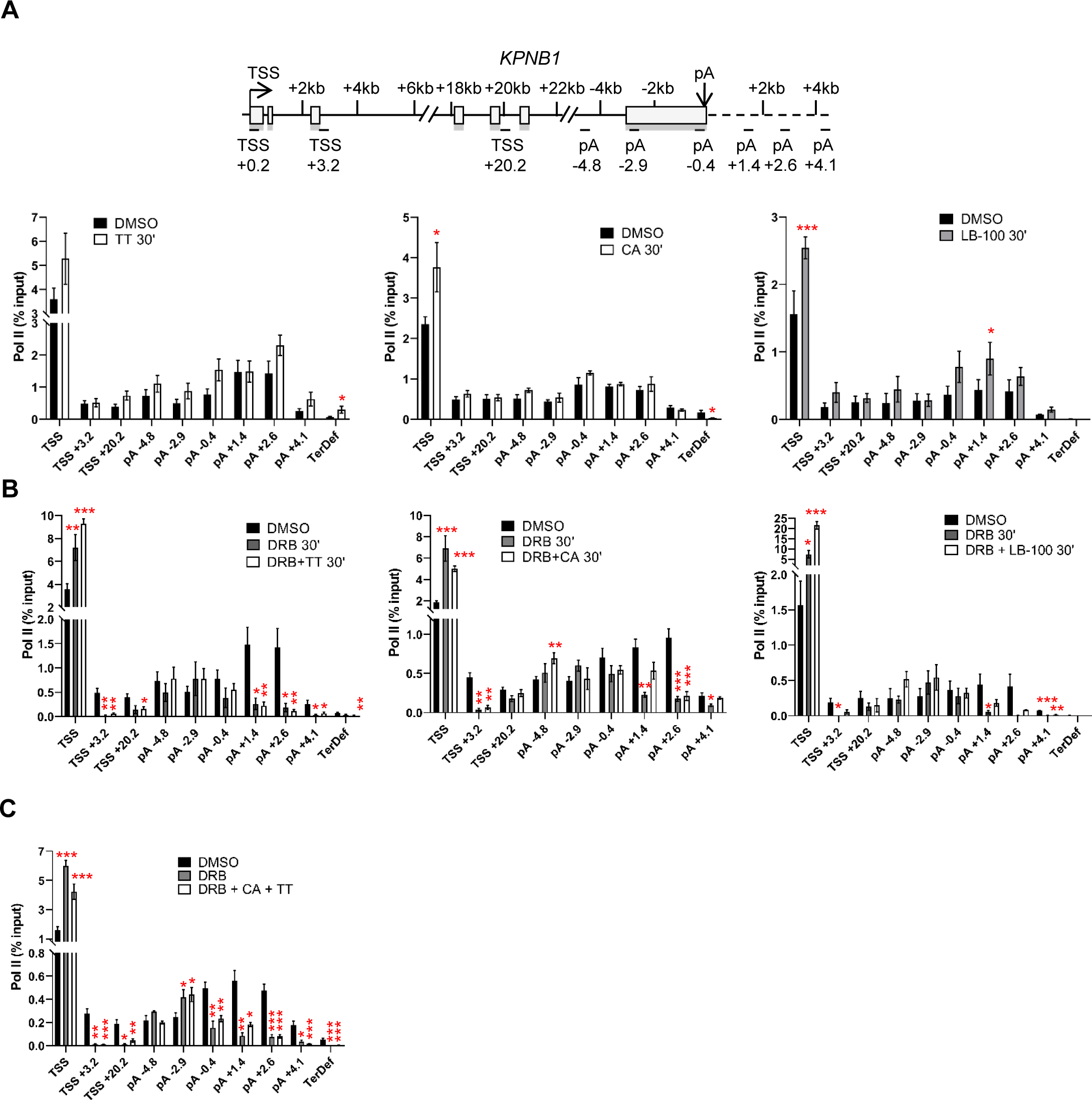
PP1 and PP2A regulate transcription of protein-coding genes. **A.** ChIP-qPCR of total pol II after 30 minutes treatment with DMSO, TT, CA, or LB- 100 on *KPNB1*. n=3 biological replicates, mean ± SEM, p-value: * p < 0.05, ** p < 0.01, *** p < 0.001. Statistical test: two-tailed unpaired t test. **B.** ChIP-qPCR of total pol II after 30 minutes treatment with DMSO, DRB, DRB + TT, DRB + CA, or DRB + LB-100 on *KPNB1*. n=3 biological replicates, mean ± SEM, p-value: * p < 0.05, ** p < 0.01, *** p < 0.001. Statistical test: two-tailed unpaired t test. **C.** ChIP-qPCR of total pol II after 30 minutes treatment with DMSO or DRB+CA+TT on the *KPNB1*. n=3 biological replicates, mean ± SEM, p-value: * p < 0.05, ** p < 0.01, *** p < 0.001. Statistical test: two-tailed unpaired t test.

To confirm our ChIP-qPCR results, we performed mouse spike-in total pol II ChIP-seq in HeLa cells treated for 30 minutes with DMSO, DRB, CA, or a combination of DRB and CA (Figure 6A- C). Inhibition of CDK9 by DRB causes an increase in pol II pausing, a loss of pol II entering productive elongation, an increase of the level of pol II across the last exon and premature termination. CA treatment results in a small increase in pol II pausing but does not affect pol II entry into productive elongation. Interestingly, inhibition of PP2A by CA promotes a specific increase in pol II across the last exon, indicating that both CDK9 and PP2A activities are required across the last exon to ensure proper transcription elongation. Inhibition of both CDK9 and PP2A at the same time does not reverse the effect of CDK9 inhibition on pol II pausing downstream from the TSS and at the EEC but abrogates the effect of CDK9 inhibition on pol II transcription at the 3’end of protein-coding genes. Thus, the combination of CDK9 and PP2A inhibition results in pol II elongating further downstream of the poly(A) site than after CDK9 inhibition alone. This is noted on two single gene examples, *SUPT5H* and *WDR43* (Figure 6D).

**Figure 6.**
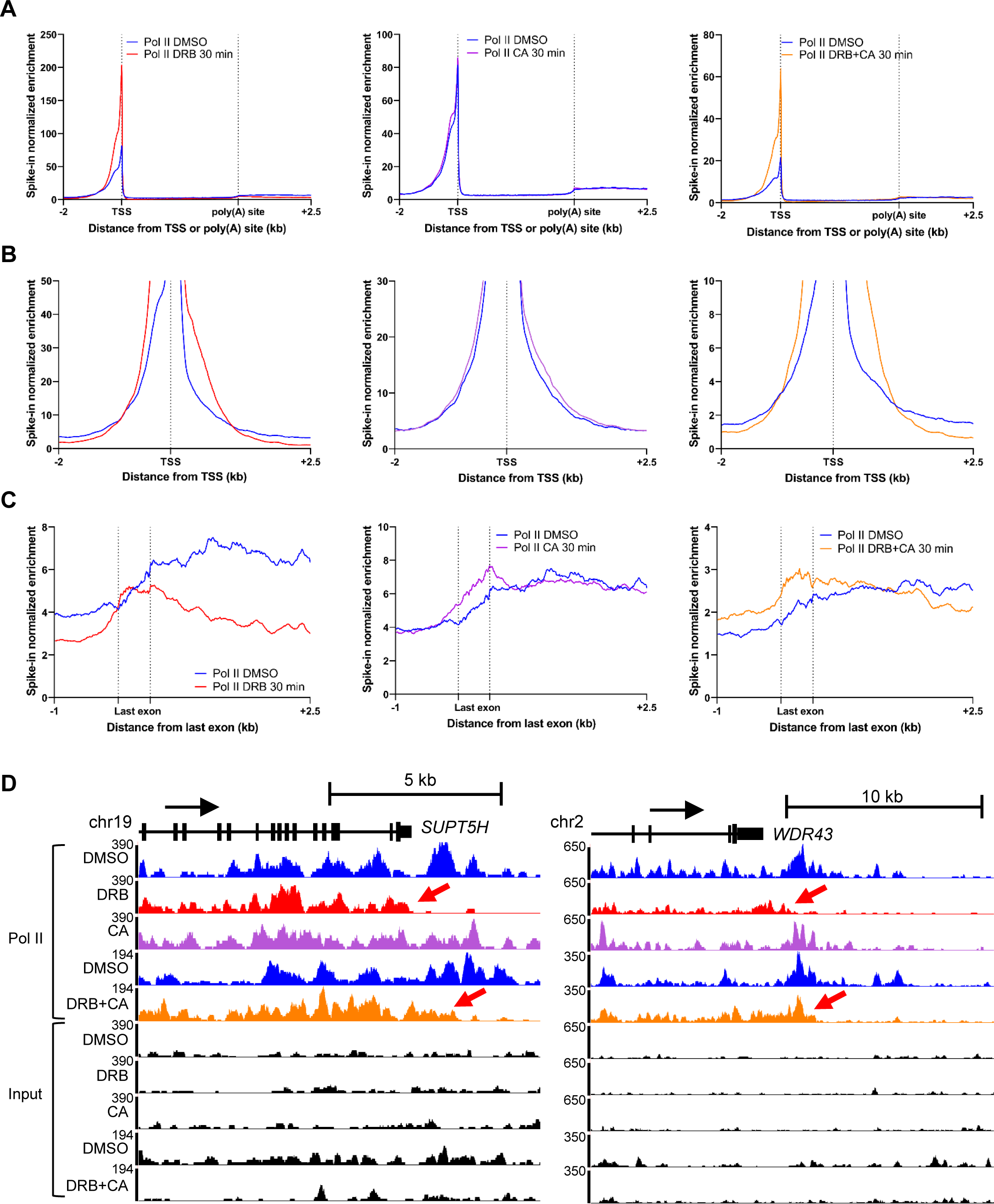
PP2A counteracts CDK9 activity at the 3’ end of genes. **A.** Metagene profile of mouse spiked-in total pol II ChIP-seq in HeLa cells after 30 minutes treatment with DMSO (blue), DRB (red), CA (purple), or DRB and CA (orange) across scaled expressed protein-coding genes longer than 30 kb (n=3,490). **B.** Metagene profile of mouse spiked-in total pol II ChIP-seq in HeLa cells after 30 minutes treatment with DMSO (blue), DRB (red), CA (purple), or DRB and CA (orange) around the TSS. Only a part of the y-axis is presented to show the entry of pol II into productive elongation. **C.** Metagene profile of mouse spiked-in total pol II ChIP-seq in HeLa cells after 30 minutes treatment with DMSO (blue), DRB (red), CA (purple), or DRB and CA (orange) around the scaled last exon of expressed genes longer than 40 kb (n=2,525). **D.** Screenshot of the genome browser of mouse spiked- in total pol II ChIP-seq tracks around the 3’end of the protein-coding genes *SUPT5H* and *WDR43*. Red arrows indicate premature termination of pol II (DRB tracks, red) and transcription further downstream the poly(A) site (DRB+CA tracks, orange).

These findings indicate that CDK9 and PP2A both play roles in regulating the progress of pol II at the 3’ end of genes.

### CDK9 and PP2A regulate mRNA cleavage and polyadenylation and alternative poly(A) site usage

As inhibition of PP2A abrogates the effect of CDK9 inhibition on transcription at the 3’end of the gene, we tested whether PP2A inhibition also reverses the effect of CDK9 inhibition on cleavage and polyadenylation. Treatment of cells for 30 minutes with DRB or DRB and CA followed by qRT-PCRs of newly-synthesized nuclear polyadenylated mRNA from TNFα- induced genes indicates that inhibiting PP2A fully reverses the effect of CDK9 inhibition on the production of polyadenylated mRNA (Figures 7A and B). Surprisingly, inhibiting PP2A alone causes increased production of polyadenylated mRNA for most of the genes tested, indicating that PP2A can act as a negative regulator of CPA. To confirm that CA does not block mRNA export, which could explain the increase in nuclear poly(A)+ mRNAs, we carried out qRT-PCR on the cytoplasmic fraction (Figure 7C). CA treatment of cells also results in an increase in poly(A)+ mRNAs in the cytoplasmic fraction, indicating that CA does not affect mRNA export. We confirmed these results with another PP2A inhibitor, LB-100 (Appendix Figure S5A-C). Inhibition of PP1 instead has little effect on polyadenylation of newly-synthesized RNA (Appendix Figure S5D and S5E).

**Figure 7.**
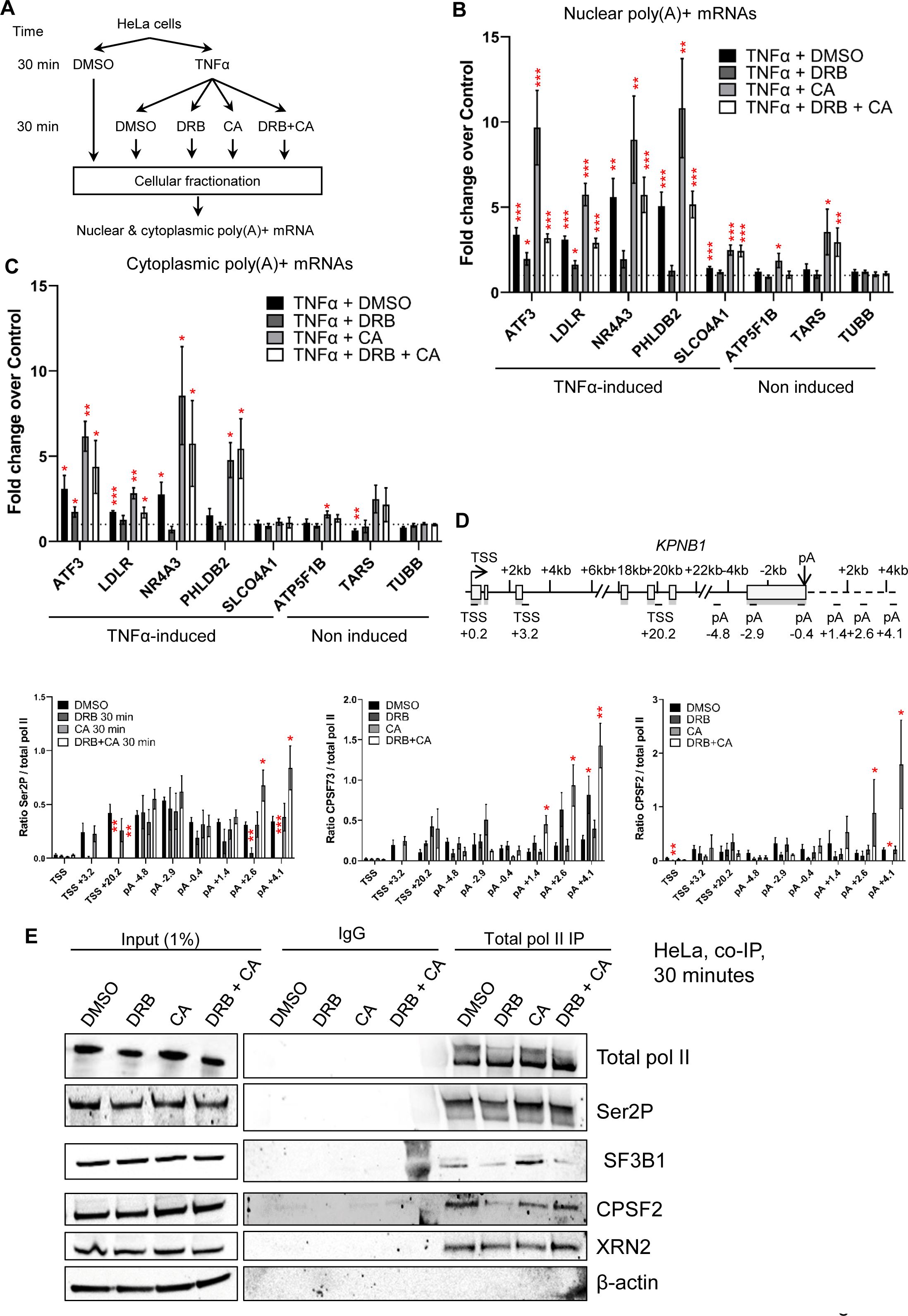
CDK9 and PP2A regulate mRNA cleavage and polyadenylation. **A.** Schematic of the nuclear and cytoplasmic qRT-PCR experiments. **B.** qRT-PCR of nuclear polyadenylated mRNAs of several TNFα induced or non-induced genes with a 30 minutes DMSO, DRB, CA, or DRB+CA treatment. n=6 biological replicates, mean ± SEM, p-value: * p < 0.05, ** p < 0.01, *** p < 0.001. Statistical test: two-tailed unpaired t test. **C.** qRT-PCR of cytoplasmic polyadenylated mRNAs of several TNFα induced or non-induced genes with a 30 minutes DMSO, DRB, CA, or DRB+CA treatment. n=4 biological replicates, mean ± SEM, p-value: * p < 0.05, ** p < 0.01, *** p < 0.001. Statistical test: two-tailed unpaired t test. **D.** ChIP-qPCR of Ser2P, CPSF73, or CPSF2 ratioed to total pol II after 30 minutes treatment with DMSO, DRB, CA, or DRB+CA on the *KPNB1*. n=3 biological replicates, mean ± SEM, p- value: * p < 0.05, ** p < 0.01, *** p < 0.001. Statistical test: two-tailed unpaired t test. **E.** Co-immunoprecipitation of total pol II from HeLa cells treated for 30 minutes with DMSO, DRB, CA, or DRB+CA followed by western blot with total pol II, Ser2P, SF3B1, CPSF2, Xrn2, and GAPDH antibodies.

To determine whether CPA factor recruitment is affected by PP2A inhibition, we performed western blots of Xrn2, CPSF2, and CPSF73 on the chromatin and nucleoplasm fractions following 30 minutes of treatment with DRB, CA, or DRB and CA (Figure EV4A-C). Whereas CDK9 inhibition causes a decrease in Xrn2, CPSF2, and CPSF73 on chromatin, CA or DRB and CA treatment causes an increase in CPSF2 and CPSF73 recruitment to chromatin and a decrease of CPSF2 in the nucleoplasmic fraction. Importantly, there was no effect of DRB, CA, or DRB and CA on the total protein level of these termination factors in whole cell extract (Figure EV4D). As pol II CTD Ser2P is associated with the recruitment of CPA factors and CDK9 and PP2A modulate Ser2P, we investigated the effect of CDK9, PP2A or CDK9/PP2A inhibition on pol II, Ser2P, CPSF73, and CPSF2 levels by ChIP-qPCR on the *KPNB1* gene. (Figure 7D and Figure EV4E). DRB and CA treatment together results in a localized increase of the Ser2P/pol II ratio and an increased ratio of CPSF73/pol II and CPSF2/pol II at the 3’end of *KPNB1*. Analysis of the nuclear polyadenylated KPNB1 mRNA level indicates that, as expected from the 3’READS experiments, it is not induced by TNFα (Figure EV4F). However, the level of nuclear polyadenylated KPNB1 mRNA increases after CA treatment while DRB and CA treatment at the same time leads to a modest increase in the poly(A)+ mRNA level compared to DRB alone. In addition, DRB reduces Ser2P while CA or DRB and CA together do not. Interestingly, DRB treatment strongly decreases the interaction between pol II and SF3B1 and between pol II and CPSF2 and to a lesser extent between pol II and Xrn2 (Figure 7E). Interestingly, CA treatment alone increases the interaction between SF3B1 and pol II while the interaction between pol II and CPSF2 and Xrn2 is not affected. In contrast, the interaction between pol II and SF3B1 is still decreased by DRB and CA treatment while the interaction between pol II and CPSF2 and Xrn2 is unaffected, indicating that CDK9 and PP2A regulate the interaction between pol II, SF3B1, and CPA complex components.

Further analysis of our 3’READS data indicates that 30 minutes of CDK9 inhibition has only a limited effect on intronic poly(A) site (IPA) usage, with only 4 genes exhibiting increased IPA and 3 genes exhibiting decreased IPA (Figure EV5A). In addition, 22 genes showed a shift towards proximal poly(A) site usage and 17 genes showed a shift towards distal poly(A) site usage following CDK9 inhibition (Figure EV5B). For three genes with a shift towards proximal poly(A) site usage after CDK9 inhibition in the 3’READS data, *EIF1*, *HCCS*, and *PCF11*, this was confirmed by qRT-PCR (Figure EV5C and EV5D). PP2A inhibition promotes a shift towards distal poly(A) site usage on *EIF1* and *HCCS* (Figure EV5D). However, inhibiting both CDK9 and PP2A together reverses most of the effect of CDK9 inhibition on the three genes, with only *PCF11* still exhibiting some increase in proximal poly(A) site usage compared to the control. These finding reinforce the role of the CDK9/PP2A kinase/phosphatase pair in regulating both poly(A) site function and transcription at the end of protein-coding genes.

## Discussion

We previously showed that CDK9 inhibition, in addition to halting pol II at the EEC, leads to premature termination of pol II close to the poly(A) site (Laitem et al., 2015). We show here that premature termination of pol II is associated with loss of mRNA polyadenylation and loss of recruitment of polyadenylation and termination factors to chromatin. Although we have termed this 3’ end CDK9 checkpoint the poly(A)-associated checkpoint, analysis of pol II transcription at single nucleotide resolution using mNET-seq indicates that pol II slows down prematurely from the start of the last exon. Thus, CDK9 inhibition may be causing failure to properly define the last exon, which helps to ensure the correct transition of pol II between elongation and termination (Figure 8). In support of this, we identified several proteins involved in definition of the last exon and the transition between transcription elongation and termination, including SPT5, SF3B1, CDC5L and the RNA m6a methyltransferase METTL3, (Cortazar et al., 2019, Ke, Alemu et al., 2015, Kyburz et al., 2006, Parua et al., 2020, Tellier et al., 2020a) as targets of CDK9. CDK9 therefore joins other transcriptional CDKs, including CDK11 and CDK12, in the regulation of mRNA CPA, emphasizing the critical role of protein phosphorylation in this co-transcriptional process (Davidson, Muniz et al., 2014, Eifler, Shao et al., 2015, Pak, Eifler et al., 2015, Tellier et al., 2020b). Surprisingly, there is little overlap in targets between all the different CDK9 phosphoproteomic experiments published (Decker et al., 2019, Sanso et al., 2016). This is likely explained by differences in the CDK9 inhibitors (DRB versus analog-sensitive cell line), *in vitro* vs *in vivo* approaches, and also different cell lines used (HeLa, Raji CDK9as).

**Figure 8.**
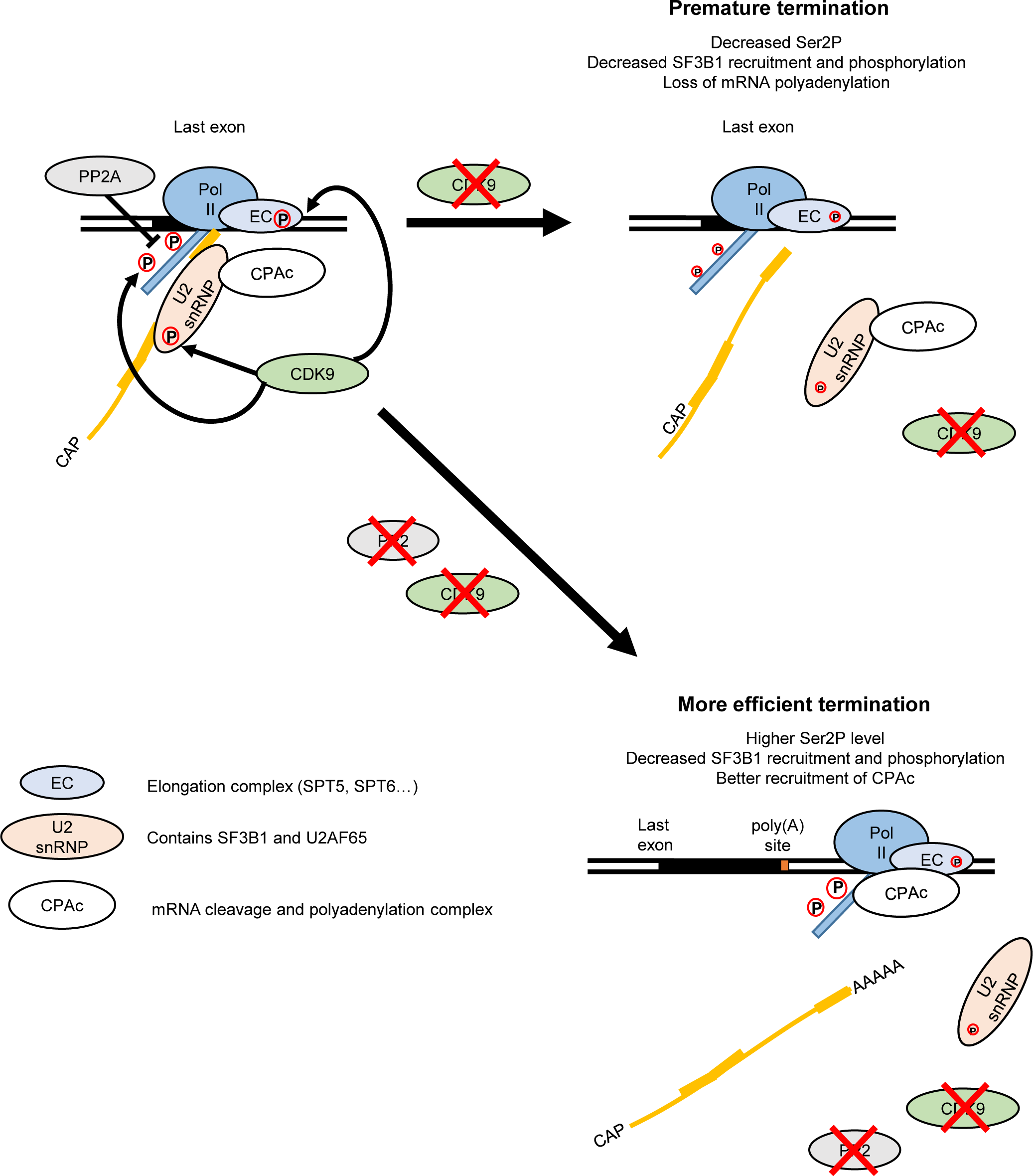
Model of CDK9 and PP2A functions in transcription termination and RNA maturation. During the transcription cycle, CDK9 phosphorylates the pol II CTD on Ser2 and Ser5, proteins found in the elongation complex eg, SPT5, and the SF3B1 subunit of the U2 snRNP. The phosphatase PP2A dephosphorylates the pol II CTD on Ser2P and Ser5P. CDK9 inhibition causes a decrease in pol II CTD phosphorylation and proteins found in the elongation complex and loss of the SF3B complex together with CPA factors from pol II, resulting in the premature termination of pol II and loss of polyadenylation. Inhibition of CDK9 and PP2A at the same time results in more phosphorylation of the pol II CTD, but not of SF3B1 T142P. However, PP2A inhibition counteracts the effect of CDK9 inhibition on transcription and mRNA CPA, likely via pol II CTD Ser2 phosphorylation, resulting in restored mRNA maturation and subsequent transcription termination.

It has previously been shown that hyperactivation of P-TEFb via the degradation of the 7SK non-coding RNA, which negatively regulates P-TEFb activity, leads to global pol II readthrough at poly(A) sites (Castelo-Branco, Amaral et al., 2013). This in line with the demonstration that inhibition of PP1, which will mimic P-TEFb hyperactivation on for example, SPT5 and the pol II CTD, also promotes transcriptional readthrough (Cortazar et al., 2019, Eaton et al., 2020, Parua et al., 2018, Parua et al., 2020). These results underline the importance of regulating P- activity at the end of genes. However, knockout of 7SK does not result in global readthrough, (Bandiera, Wagner et al., 2021, Studniarek, Tellier et al., 2021) indicating that more work is required to understand the role this non-coding RNA plays in the process of transcription termination.

In line with previous findings, we found that CDK9 activity is needed for CTD phosphorylation on Ser2 and Ser5 *in vivo* (Czudnochowski, Bosken et al., 2012, Ghamari, van de Corput et al., 2013, Greifenberg, Honig et al., 2016, Laitem et al., 2015). As Ser2 phosphorylation helps to recruit polyadenylation factors (Davidson et al., 2014, Eifler et al., 2015, Tellier et al., 2020b), loss of this mark would be expected to affect recruitment of polyadenylation factors.

However, recruitment of the key polyadenylation factor, PCF11, which binds directly to Ser2P is not greatly affected (Laitem et al., 2015), raising the question of why the other polyadenylation factors are lost.

In addition to the previously-demonstrated roles of PP1 and PP2A in transcription regulation, at the 5’ and 3’ ends of genes, respectively (Cortazar et al., 2019, Eaton et al., 2020, Huang et al., 2020, Vervoort et al., 2021, Zheng et al., 2020), we found that inhibiting PP2A abrogates the premature termination caused by CDK9 inhibition. In addition, inhibition of PP2A reverses the loss of CTD phosphorylation and loss of CPA factors from chromatin caused by CDK9 inhibition and production of polyadenylated mRNA is restored. In contrast, inhibition of CDK9 and PP2A together does not reverse the loss of SF3B1 recruitment to the pol II caused by CDK9 inhibition, indicating that another pathway, potentially Ser2 phosphorylation, is important for recruiting the CPA complex. The CDK9/PP2A kinase/phosphatase pair is therefore involved in regulating the recruitment and activity of the CPA complex.

It has been previously shown that loss of the INTAC complex, which is composed of the PP2A core enzyme and the RNA endonuclease Integrator complex, could reverse the effect of CDK9 inhibition at the EEC (Fianu, Chen et al., 2021, Hu, Peng et al., 2021, Vervoort et al., 2021, Zheng et al., 2020). Interestingly, we did not observe reversal of the effect of CDK9 inhibition at the EEC when PP2A was inhibited by Calyculin A or LB-100, indicating that INTAC function at the EEC is independent of PP2A activity.

Although a low concentration of Tautomycetin or Calyculin A can specifically inhibit PP1 or PP2A, respectively, we cannot rule out that these small molecule inhibitors affect other proteins. However, another PP2A inhibitor, LB-100 gave similar results. In addition, PP1 and PP2A are clearly not completely redundant as PP1, but not PP2A, dephosphorylates SF3B1 T142P. In addition, both phosphatases are active at the 5’ and 3’ ends of protein-coding genes but have different functions. PP1 inhibition leads to a termination defect, likely due to a high pol II elongation rate after the poly(A) site caused by hyperphosphorylation of SPT5, which impedes Xrn2-mediated transcription termination (Cortazar et al., 2019, Eaton et al., 2020). Conversely, PP2A inhibition seems to promote more efficient cleavage and polyadenylation of transcripts. We found that PP2A inhibition promotes higher interaction between SF3B1 and total pol II while the Ser2P level or CPA factor recruitment to the pol II are not higher than in untreated conditions. A higher level of SF3B1 on the pol II complex could therefore improve the activity of the CPA complex rather than recruiting higher level of CPA factors to the pol II.

Interestingly, in addition to phospho-SPT5, phospho-SF3B1 is a target of PP1, supporting a role for this phosphatase in splicing and potentially in the definition of the last exon (Mermoud et al., 1992, Shi et al., 2006). However, inhibition of PP1 does not reverse the loss of Ser2 and Ser5 phosphorylation caused by CDK9 inhibition. This result reinforces the notion that dephosphorylation of Ser2P and Ser5P by PP1 is redundant with other CTD phosphatases, including PP2A.

Our DRB mNET-seq time course indicates that after 5 minutes of CDK9 inhibition, the effect on pol II at the EEC is more drastic than at the 3’end of the genes, where a defect becomes clear only after 10 minutes of inhibition. The drastic effect of CDK9 inhibition on mRNA CPA may indicate that the loss of mRNA polyadenylation precedes premature termination of pol II. Thus, dephosphorylation of CDK9 targets including the pol II CTD, SPT5, and SF3B1 and the loss of the CPA complex from chromatin would occur first, and indeed, the loss of CPA factors is stronger than the loss of pol II after 30 minutes of CDK9 inhibition (Figures 1F and 7D). The slowing down of pol II over the last exon and early disengagement of pol II may be caused by the subsequent loss of elongation factors, such as SPT5, SPT6, or the PAF1 complex in turn, as these keep pol II clamped onto the DNA template (Bernecky, Plitzko et al., 2017, Hou, Wang et al., 2019, Vos et al., 2018a). Loss of the CPA complex is likely to be linked to loss of pol II CTD phosphorylation, which helps to recruit CPA factors to pol II and/or loss of an SF3B/CPA complex. Reversal of this by PP2A inhibition may be effected by the increase in CTD/SF3B phosphorylation followed by recovery of the normal pol II elongation rate/processivity and efficient co-transcriptional processes (pre-mRNA splicing and mRNA cleavage and polyadenylation). Currently, we cannot rule out any of these factors.

A decrease in the pol II level downstream of the poly(A) site can therefore be associated with premature non-productive termination, where gene expression is aborted (CDK9 inhibition) or more efficient termination coupled to the production of mature polyadenylated mRNA (CDK9 and PP2A inhibition) (Figure 8). Premature termination of pol II caused by inhibition of CDK9 is associated with a decreased level of Ser2P and Ser5P and reduced recruitment of CPA factors. In contrast, inhibition of CDK9 and PP2A together promote more efficient termination of pol II with the production of de novo nuclear polyadenylated mRNA, which is associated with higher Ser2P, efficient recruitment of CPA factors, and termination closer to the poly(A) site.

## Materials and methods

### Cell culture

HEK293 and HeLa cells were obtained from ATCC (ATCC® CRL-1573™ and ATCC® CCL-2™, respectively). HeLa, HEK293 parental cells, and CDK9as HEK293 cells were grown in DMEM medium supplemented with 10% foetal calf serum, 100 U/ml penicillin, 100 μg/ml streptomycin, 2 mM L-glutamine at 37°C and 5% CO2. HEK293 and CDK9as cells were treated with 7.5, 10, or 15 μM 1-NA-PP1 (Cayman Chemical Company) for 15 and 30 minutes. HEK293, CDK9as, or HeLa cells were treated with 10 ng/ml of TNFα (PeproTech), 2.5 nM Calyculin A (Sigma), 25 nM Tautomycetin (Bio-Techne), 2.5 nM LB-100 (Stratech Scientific Ltd), 100 μM DRB (Sigma) for 5, 10, 15 or 30 minutes, or 0.5 or 1 μM SNS-032 (LKT labs). As a negative control, HEK293, CDK9as, and HeLa cells were treated with DMSO (the resuspension vehicle for NA, DRB, Calyculin A, LB-100, and Tautomycetin). Cells were routinely checked to be free of mycoplasma contamination using Plasmo Test Mycoplasma Detection Kit (InvivoGen, rep- pt1).

### Analog sensitive cell line creation

Guide RNAs were computationally designed.

Guide RNA 1: 5’-GCTCGCAGAAGTCGAACACC-3’

Guide RNA 2: 5’-CTTCTGCGAGCATGACCTTGC-3’

The modified CDK9 genomic sequence (NCBI RefSeq Accession NG_033942.1) (500 bp either site of the mutation) was cloned into pcDNA3 and used as the repair template for genome editing. The repair template contains a TTC (phenylalanine) to GCT (alanine) mutation. The guide (g)RNAs inserts were cloned into the pX462 vector (obtained from Addgene). HEK293 cells were transfected with the gRNA vectors and correction template using Lipofectamine 2000 (Life Technologies) following the manufacturer instructions. Single clones were isolated by low density plating after Puromycin and Neomycin selection. Genomic DNA from each clone was analysed using PCR and Sanger sequencing.

### gDNA preparation

HEL293 and CDK9as cells were incubated in 180 μl ChIP lysis buffer (10 mM Tris–HCl ph8.0, 0.25% Triton X-100, 10 mM EDTA) for 10 min and sonicated for 3 min (30 seconds on/30 seconds off) using a Q800R2 sonicator (QSONICA). 20 μl ammonium acetate was added (final concentration of 400 mM) and samples were mixed with 200 μl phenol/chloroform by vortexing. The samples were centrifuged at 13,000 g for 5 minutes at 25°C and the upper phase was transferred to a new tube. Genomic DNA was precipitated in 70% ethanol, pelleted by centrifugation at 13,000 g for 5 minutes at 25°C and dissolved in nuclease free water. A CDK9 fragment was PCR amplified using the following primers: Forward: 5′- AAGGCTTCTGAGACAGCTGG-3′; Reverse: 5′-CAACCAGCTTCTTTCTTCCTGC-3′. DNA was purified using a QIAquick PCR purification kit (Qiagen) and sequenced by the Source Bioscience Sanger Sequencing Service, Oxford.

### RNA preparation

RNA was extracted from HEK293 and CDK9as cells using a *Quick*-RNA Miniprep kit (Zymo Research) according to the manufacturer’s instructions. Reverse-transcription (RT) was performed with 500 ng of RNA using random hexamers with the SuperScript III kit (Invitrogen) according to the manufacturer’s instructions. Sequencing was performed on a cDNA PCR fragment generated with Forward primer: 5’-AAAGCAGTACGACTCGGTGG-3’ and Reverse primer: 5’-GTAGAGGCCGTTAAGCAGCA-3’, purified using a QIAquick PCR purification kit (Qiagen) and sequenced by the Source Bioscience Sanger Sequencing Service, Oxford.

### Cell proliferation analysis

Cells were seeded at 500/well in 95 μl in 96-well microplates (Greiner, 655090) and measured every 12 or 24 hours by adding alamarBlue HS (Invitrogen) (1/20) and reading in a fluorimeter after 1 hour’s incubation, according to the manufacturer’s instructions. 1-NA-PP1 or DMSO was added to the cells as noted on the figure.

### Chromatin immunoprecipitation (ChIP)

ChIP analysis were performed as previously described (Tellier et al., 2020b). HeLa, HEK293, and CDK9as cells were grown in 100 or 150 mm dishes until they reached ∼80% confluency. The cells were fixed with 1% formaldehyde for 10 minutes at room temperature with shaking. Formaldehyde was quenched with 125 mM glycine for 5 minutes at room temperature with shaking. The cells were washed twice with ice-cold PBS, scraped with ice-cold PBS, and transferred into 1.5 or 15 ml Eppendorf tubes. Cells were pelleted for 10 minutes at 1,500 rpm at 4°C. The pellets were then resuspended in ChIP lysis buffer (10 mM Tris–HCl ph8.0, 0.25% Triton X-100, 10 mM EDTA, protease inhibitor cocktail, and phosphatase inhibitor) and incubated 10 minutes on ice before being centrifuged at 1,500 g for 5 minutes at 4°C. Pellets were resuspended in ChIP wash buffer (10 mM Tris–HCl pH8.0, 200 mM NaCl, 1 mM EDTA, protease inhibitor cocktail, and phosphatase inhibitor) and centrifuged at 1,500 g for 5 minutes at 4°C. Pellets were resuspended in ChIP sonication buffer (10 mM Tris–HCl pH 8.0, 100 mM NaCl, 1 mM EDTA, protease inhibitor cocktail, and phosphatase inhibitor) and incubated 10 minutes on ice. HeLa cells were sonicated for 30 cycles, 30 seconds on/30 seconds off using a Bioruptor Pico (Diagenode). HEK293 and CDK9as cells were sonicated for one hour, 30 seconds on/30 seconds off, 40% amplitude, using a Q800R2 sonicator (QSONICA). Chromatin was pelleted at 13,000 rpm for 15 minutes at 4°C and supernatant transferred to a new Eppendorf tube.

Chromatin was pre-cleared for 30 min on a rotating wheel at 4°C with 10 µl of Protein G Dynabeads, previously washed with 100 µl of RIPA buffer (10 mM Tris–HCl pH8.0, 150 mM NaCl, 1 mM EDTA, 0.1% SDS, 1% Triton X-100, 0.1% sodium deoxycholate). Chromatin was quantified on a NanoDrop One with 60-100 µg of chromatin used per IP (antibodies described in Appendix Table 1) and incubated overnight on a rotating wheel at 4°C. 15 μl of Dynabeads per IP were washed in 100 μl RIPA buffer. The beads were saturated with 15 μl RIPA containing 4mg/ml of bovine serum albumin (BSA) and mixed overnight on a rotor at 4°C.

Dynabeads were then mixed for 1 hour on a rotating wheel at 4°C with the chromatin incubated with the antibody. Beads were then washed three times with 300 μl ice-cold RIPA buffer, three times with 300 μl High Salt Wash buffer (10 mM Tris–HCl pH8.0, 500 mM NaCl, 1 mM EDTA, 0.1% SDS, 1% Triton X-100, 0.1% sodium deoxycholate), twice with 300 μl LiCl Wash buffer (10 mM Tris–HCl pH8.0, 250 mM LiCl, 1 mM EDTA, 1% NP-40, 1% sodium deoxycholate), and twice with 300 μl TE buffer (10 mM Tris–HCl pH 7.5, 1 mM EDTA). Each sample was eluted twice from the Dynabeads with 50 μl of Elution buffer (100 mM NaHCO3, 1% SDS, 10 mM DTT) for 15 minutes at 25°C at 1,400 rpm on a Thermomixer. For each input sample, 90 μl of Elution buffer was added to 10 μl total input. Each sample was treated with RNase A (0.6 μl of 10 mg/ml) for 30 minutes at 37°C followed by the addition of 200 mM NaCl and a five hours incubation at 65°C to reverse the crosslinks. Precipitation was performed overnight at -20°C following the addition of 2.5x volume of 100% ethanol. Ethanol was removed after a 20 minutes centrifugation at 13,000 rpm at 4°C and pellets resuspended in 100 μl TE, 25 μl 5x Proteinase K buffer (50 mM Tris–HCl pH 7.5, 25 mM EDTA, 1.25% SDS) and 1.5 μl Proteinase K (20 mg/ml). The samples were incubated two hours at 45°C to degrade the proteins. DNA was purified using Qiagen PCR Purification Kit and kept at -20°C.

ChIP samples were analysed by real-time qPCR using QuantiTect SYBR Green PCR kit (Qiagen) and Rotor-Gene RG-3000 (Corbett Research). Signals are presented as percentage of Input after removing the background signal from the IP with the IgG antibody. The sequence of primers used for ChIP-qPCR is given in Appendix Table 2. Experiments were replicated three times and each ChIP sample was measured in triplicate by qPCR.

ChIP-seq of total pol II (Active Motif, 39097) in HeLa cells treated with DMSO, 100 μM DRB, 2.5 nM CA, or 100 μM DRB + 2.5 nM CA were performed in biological duplicates. Preparation of ChIP-seq library and ChIP sequencing was prepared with the NEBNext Ultra II DNA Library Prep Kit for Illumina (NEB), according to the manufacturer’s instructions. DNA sequencing was conducted by the high throughput genomics team of the Wellcome Trust Centre for Human Genetics (WTCHG), Oxford or by Novogene UK.

### Co-immunoprecipitation

For each sample and IgG control: 120 μl of Dynabeads M-280 Sheep anti-mouse IgG (Thermo Fisher) or 40 μl of Dynabeads protein G (Thermo Fisher) were pre-blocked overnight at 4°C on a wheel in 1 ml of PBS supplemented with 0.5% BSA. The next day, the beads were washed three times in IP buffer (25 mM Tris–HCl pH 8.0, 150 mM NaCl, 0.5% NP-40, 10% Glycerol, 2.5 mM MgCl2), before being incubated for two hours at 4°C on a wheel in 600 μl of IP buffer supplemented with 5 μg of total pol II antibody (MABI0601, MBL International), 5 μg of SF3B1 antibody (D221-3, MBL International), 5 μg of CPSF2 antibody (A301-581A, Bethyl Laboratories), or Normal Rabbit IgG (2729S, Cell Signaling Technology) and protease inhibitor cocktail (cOmplete™, EDTA-free Protease Inhibitor Cocktail, Sigma-Aldrich). In the meantime, a 70–80% confluent 15 cm dish of HeLa cells was washed twice with ice-cold PBS and scrapped with ice-cold PBS supplemented with protease inhibitor cocktail. The cells were pelleted at 500 g for 5 minutes at 4°C. The pellets were re-suspended in 800 μl of Lysis buffer (50 mM Tris–HCl pH 8.0, 150 mM NaCl, 1% NP-40, 10% glycerol, 2.5 mM MgCl2, protease inhibitor cocktail, PhosSTOP (Sigma-Aldrich), 1× PMSF (Sigma-Aldrich), and 25–29 units of Benzonase (Merck Millipore)) and incubated at 4°C on a wheel at 16 rpm for 30 minutes. After centrifuging for 15 minutes at 13,000 g at 4°C, 800 μl of Dilution buffer (150 mM NaCl, 10% glycerol, 2.5 mM MgCl2, protease inhibitor cocktail, PhosSTOP, and 1× PMSF) was added to each supernatant The beads conjugated with antibodies were washed three times with IP buffer supplemented with protease inhibitor cocktail before being incubated with 1 mg of proteins at 4°C on a wheel at 16 rpm for 2 hours. The beads were washed three times with IP buffer supplemented with protease inhibitor cocktail and three times with IP buffer without NP-40 supplemented with protease inhibitor cocktail. Proteins were eluted in 40 μl of 1× LDS plus 100 mM DTT for 10 minutes at 70°C. Western blots were performed with NuPAGE Novex 3–8% Tris-Acetate Protein Gels (Life Technologies). For the SF3B1 immunoprecipitation followed by proteomics, glycerol was removed from each buffer.

### Protein extraction and western blot

Western blot analysis was performed on chromatin and nucleoplasm extracts as previously described in the mNET-seq procedure (Nojima, Gomes et al., 2015). A ∼80% confluent 15 cm dish was washed twice with ice-cold PBS and scrapped in 5 ml of ice-cold PBS. The cells were pelleted at 420 g for 5 minutes at 4°C. After discarding the supernatant, the cells were resuspended in 4 ml of ice-cold HLB+N buffer (10mM Tris-HCl (pH 7.5), 10 mM NaCl, 2.5 mM MgCl2 and 0.5% (vol/vol) NP-40) and incubated on ice for 5 minutes. The cell pellets were then underlayed with 1 ml of ice-cold HLB+NS buffer (10 mM Tris-HCl (pH 7.5), 10 mM NaCl, 2.5 mM MgCl2, 0.5% (vol/vol) NP-40 and 10% (wt/vol) sucrose). Following centrifugation at 420 g for 5 minutes at 4°C, the nuclear pellets were resuspended by pipetting up and down in 125 μl of NUN1 buffer (20 mM Tris-HCl (pH 7.9), 75 mM NaCl, 0.5 mM EDTA and 50% (vol/vol) glycerol) and moved to a new 1.5 ml ice-cold microcentrifuge tube. Following the addition of 1.2 ml of ice-cold NUN2 buffer (20 mM HEPES-KOH (pH 7.6), 300 mM NaCl, 0.2 mM EDTA, 7.5 mM MgCl2, 1% (vol/vol) NP-40 and 1 M urea), the tubes were vortexed at maximum speed for 10 seconds and incubated on ice for 15 minutes with a vortexing step of 10 seconds every 3 minutes. The samples were centrifuged at 16,000 g for 10 minutes at 4°C and the supernatant kept as the nucleoplasmic fractions while the chromatin pellets were washed with 500 μl of ice-cold PBS and then with 100 μl of ice-cold water. The chromatin pellet was then digested in 100 μl of water supplemented with 1 μl of Benzonase (25–29 units, Merck Millipore) for 15 minutes at 37°C in a thermomixer at 1,400 rpm. 10 μg of proteins were boiled in 1× LDS plus 100 mM DTT. Western blots were performed with NuPAGE Novex 4–12% Bis–Tris Protein Gels (Life Technologies).

For whole cell extract, cells were washed in ice-cold PBS twice, collected in ice-cold PBS with a 3,000 rpm centrifugation for 5 minutes at 4°C. The pellets were re-suspended in RIPA buffer supplemented with protease inhibitor cocktail and PhosSTOP, kept on ice for 30 minutes with a vortexing step every 10 minutes. After centrifugation at 14,000 g for 15 minutes at 4°C, the supernatants were kept and quantified with the Bradford method. 20 μg of proteins were boiled in 1× LDS plus 100 mM DTT. Western blots were performed with NuPAGE Novex 4–12% Bis–Tris Protein Gels (Life Technologies). The list of primary antibodies is shown in Appendix Table 1.

Secondary antibodies were purchased from Merck Millipore (Goat Anti-Rabbit IgG Antibody, HRP-conjugate, 12-348, and Goat Anti-Mouse IgG Antibody, HRP conjugate, 12-349), the chemiluminescent substrate (SuperSignal West Pico PLUS) from Thermo Fisher, and the membranes visualized on an iBright FL1000 Imaging System (Thermo Fisher). Quantification of the western blots was performed with Image Studio Lite software.

### RNA subcellular fractionation

RNA subcellular fractionation as performed as described before (Neve et al., 2016). A ∼80% confluent 10 or 15 cm dish was washed twice with ice-cold PBS and scrapped in ice-cold PBS. The cells were pelleted at 1,000 rpm for 5 minutes at 4°C and then resuspended with slow pipetting in 1 ml of Lysis Buffer B (10 mM Tris-HCl pH 8, 140 mM NaCl, 1.5 mM MgCl2, 0.5 % NP-40). Following centrifugation at 1,000 g for 3 minutes at 4°C, 500 μl of the supernatant was moved to a new tube, centrifuged at 10,000 g for one minute at 4°C, and the supernatant moved to a new tube as the purified cytoplasmic fraction. The pellets were resuspended in 1 ml of Lysis Buffer B and 100 μl of the Detergent Stock Solution (3.3 % (w/v) sodium deoxycholate, 6.6 % (v/v) Tween 40) was added under slow vortexing. Following a 5 minutes incubation on ice, the nuclei were spun down at 1,000 g for 3 minutes at 4°C. The nuclei pellet was then washed once more in 1 ml of Lysis Buffer B and spun down at 1000 g for 3 minutes at 4°C. The nuclei pellet was resuspended in 1 ml of TRIzol using a 21-gauge syringe while 500 μl of TRIzol was added to the cytoplasmic fraction and incubated 5 minutes at room temperature. Following the addition of 100 μl or 200 μl of chloroform to the cytoplasmic or nuclear fraction, respectively, the samples were vortexed vigorously for 15 seconds and spun at 12,000 g for 15 minutes at 4°C. The aqueous fraction was transferred to a new tube containing 580 μl (nuclear) or 750 μl (cytoplasmic) of isopropanol. After a 10 minutes incubation at room temperature, the samples were spun at 12,000 g for 10 minutes at 4°C. The pellets were resuspended in 87 μl of water, 10 μl of 10 X DNase buffer, 2 μl of DNase I (Roche), and 1 μl of RNase OUT (ThermoFisher Scientific), and incubated 30 minutes at 32°C. RNA were then purified twice with phenol:chloroform pH 4.5 extraction, precipitated overnight in ethanol, and finally resuspended in nuclease free water and concentrations determined using a NanoDrop One.

### qRT-PCR

For each qRT-PCR reaction, 500 ng of RNA were reverse transcribed with Oligo(dT)12-18 Primer (ThermoFisher Scientific) and the SuperScript III kit (ThermoFisher Scientific), according to the manufacturer’s instructions. cDNA was amplified by qPCR with a QuantiTect SYBR Green PCR kit (QIAGEN) and a Rotor-Gene RG-3000 (Corbett Research). The sequence of primers used for qRT-PCR is given in Appendix Table 2. Values are normalized to GAPDH mRNA, used as control. Experiments were replicated at least three times to ensure reproducibility, and each RNA sample was measured in triplicate by qPCR.

### 3’READS Protocol

The 3’READS protocol was originally described in (Hoque, Ji et al., 2013). Briefly, 25-30 μg of RNA was subjected to one round of poly(A) selection using the Poly(A)PuristTM MAG kit (Ambion) according to the manufacturer’s protocol, followed by fragmentation using Ambion’s RNA fragmentation kit at 70°C for 5 minutes. Poly(A)-containing RNA fragments were isolated using the CU5T45 oligo (a chimeric oligo containing 5 Us and 45 Ts, Sigma) which were bound to the MyOne streptavidin C1 beads (Invitrogen) through biotin at its 5’ end. Binding of RNA with CU5T45 oligo-coated beads was carried out at room temperature for 1 hour in 1x binding buffer (10 mM Tris-HCl pH 7.5, 150 mM NaCl, 1 mM EDTA), followed by washing with a low salt buffer (10 mM Tris-HCl pH 7.5, 1 mM NaCl, 1 mM EDTA, 10% formamide). RNA bound to the CU5T45 oligo was digested with RNase H (5U in 50 µl reaction volume) at 37°C for 1 hour, which also eluted RNA from the beads. Eluted RNA fragments were purified by phenol:chloroform extraction and ethanol precipitation, followed by phosphorylation of the 5’ end with T4 kinase (NEB). Phosphorylated RNA was then purified by the RNeasy kit (Qiagen) and was sequentially ligated to a 5’-adenylated 3’-adapter (5’- rApp/NNNNGATCGTCGGACTGTAGAACTCTGAAC/3ddC) with the truncated T4 RNA ligase II (Bioo Scientific) and to a 5’ adapter (5’- GUUCAGAGUUCUACAGUCCGACGAUC) with T4 RNA ligase I (NEB). The resultant RNA was reverse-transcribed to cDNA with Superscript III (Invitrogen) followed by a library preparation with the NEBNext Fast DNA Library Prep Set for Ion Torrent (NEB). cDNA libraries were sequenced on an Ion Torrent Proton.

### mNET-seq and library preparation

mNET-seq was carried out as previously described (Nojima et al., 2015) with minor changes. In brief, the chromatin fraction was isolated from four ∼80% confluent 15 cm dish cells treated with DMSO or DRB (5, 10, 15, or 30 minutes). Chromatin was digested in 100 μl of MNase (40 units/μl) reaction buffer for 2 minutes at 1,400 rpm at 37°C in a Thermomixer. MNase was inactivated by the addition of 10 μl EGTA (25mM). The soluble digested chromatin was collected after centrifugation at 13,000 rpm for 5 minutes at 4°C. The supernatant was diluted with 400 μl of NET-2 buffer and antibody-conjugated Dynabeads M-280 Sheep anti-mouse IgG (ThermoFisher Scientific) beads were added. Antibodies used: Pol II (MABI0601, MBL International) and Ser5P (MABI0603, MBL International). Immunoprecipitation was performed at 4°C for one hour. The beads were washed in the cold room six times with 1 ml of NET-2 buffer, and once with 100 μl of 1xPNKT (1xPNK buffer and 0.05% Triton X-100) buffer. Washed beads were incubated in 200 μl PNK reaction mix at 1,400 rpm at 37°C in a Thermomixer for 6 minutes. After the reaction, beads were washed once with 1 ml of NET-2 buffer and RNA was extracted with Trizol reagent. RNA was suspended in urea Dye and resolved on 6% TBU gel (ThermoFisher Scientific) at 200 V for 5 minutes. In order to size select 35–100 nt RNAs, a gel fragment was cut between BPB and XC dye markers. A 0.5 ml tube was prepared with 3–4 small holes made with 25G needle and placed in a 1.5 ml tube. Gel fragments were placed in the layered tube and broken down by centrifugation at 12,000 rpm for 1 minute at room temperature. The small RNAs were eluted from the gel using RNA elution buffer (1 M NaOAc and 1 mM EDTA) at 25°C for one hour on a rotating wheel at 16 rpm at room temperature. Eluted RNA was purified with SpinX column (Coster) with two glass filters (Millipore) and the flow-through RNA was ethanol precipitated. RNA libraries were prepared according to manual of TruSeq Small RNA Library Preparation Kit (Illumina). 12–14 cycles of PCR were used to amplify the library. Libraries were resolved on a 6% TBE polyacrylamide gel (ThermoFisher Scientific), size-selected to remove primer-primer ligated DNA, and eluted from the gel with the RNA elution buffer. Deep sequencing (Hiseq4000, Illumina) was conducted by the high throughput genomics team of the Wellcome Trust Centre for Human Genetics (WTCHG), Oxford.

### SF3B1 proteomics

#### Sample Processing Protocol

SF3B1 duplicate pull-down samples were either digested in-solution after glycine elution at pH 2.3 and neutralization, or digested on-beads. On-beads and In-solution protein samples were respectively denaturated with 8M or 4M urea in ammonium bicarbonate buffer (100 mM) for 10 minutes at room temperature. After denaturation, cysteines were reduced with of TCEP (10 mM) for 30 minutes at room temperature and alkylated with 2-Chloroacetamide (50 mM) for 30 minutes at room temperature in the dark. Samples were then pre-digested with LysC (1 µg/100 µg of sample) for 2 hours at 37°C. Before overnight digestion with trypsin (1 µg/40 µg of sample) at 37°C, urea was diluted down to 2M in ammonium bicarbonate buffer (100 mM) and calcium chloride was added at 2mM final. The next day, tryptic digestion was stopped with the addition of formic acid (5%). Digested peptides were centrifuged for 30 minutes at 13,200rpm at 4°C to remove undigested material. Supernatant was loaded onto handmade C18 stage tip, pre-activated with 100% acetonitrile, by centrifugation at 4,000rpm at room temperature. Peptides were washed twice in TFA 0.1%, eluted in 50% acetonitrile /0.1% TFA and speed-vacuum dried. Peptides were resuspended into 2% acetonitrile / 0.1% formic acid before LC-MS/MS analysis. Peptides were separated by nano liquid chromatography (Thermo Scientific Easy-nLC 1000) coupled in line a Q Exactive mass spectrometer equipped with an Easy-Spray source (Thermo Fischer Scientific). Peptides were trapped onto a C18 PepMac100 precolumn (300µm i.d.x5mm, 100Å, ThermoFischer Scientific) using Solvent A (0.1% Formic acid, HPLC grade water). The peptides were further separated onto an Easy-Spray RSLC C18 column (75um i.d., 50cm length, Thermo Fischer Scientific) using a 60 minutes linear gradient (15% to 35% solvent B (0.1% formic acid in acetonitrile)) at a flow rate 200nl/min. The raw data were acquired on the mass spectrometer in a data-dependent acquisition mode (DDA). Full-scan MS spectra were acquired in the Orbitrap (Scan range 350-1500m/z, resolution 70,000; AGC target, 3e6, maximum injection time, 100ms). The 10 most intense peaks were selected for higher-energy collision dissociation (HCD) fragmentation at 30% of normalized collision energy. HCD spectra were acquired in the Orbitrap at resolution 17,500, AGC target 5e4, maximum injection time 120ms with fixed mass at 180m/z. Charge exclusion was selected for unassigned and 1+ ions. The dynamic exclusion was set to 20 s.

#### Data Processing Protocol

Tandem mass spectra were searched using Sequest HT in Proteome discoverer software version 1.4 against a protein sequence database containing 20,405 protein entries, including 20,122 Homo sapiens proteins (Uniprot database release of 2020-04) and 283 common contaminants. During database searching cysteines (C) were considered to be fully carbamidomethylated (+57,0215, statically added), methionine (M) to be fully oxidised (+15,9949, dynamically added), all N-terminal residues to be acetylated (+42,0106, dynamically added). Two missed cleavages were permitted. Peptide mass tolerance was set at 50ppm on the precursor and 0.6 Da on the fragment ions. Data was filtered at FDR below 1% at PSM level.

### SILAC phosphoproteomics

SILAC phosphoproteomics was performed as previously described (Poss, Ebmeier et al., 2016). For stable isotope labelling with amino acids in cell culture (SILAC), Hela cells were grown in DMEM media for SILAC (minus L-Lysine and L-Arginine, Fisher Scientific) and with SILAC dialysed Foetal Bovine Serum (Dundee Cell Products). The medium was supplemented with either Arg10 (33.6 mg/ml) and Lys8 (73 mg/ml) or Arg0 and Lys0 for heavy and light treatment, respectively. After six passages at 1:3 ratio, SILAC incorporation test in HeLa cells was validated by mass spectrometry analysis.

Cells were passaged 7-8 times in SILAC media on 15 cm dishes. For each replicate, approximately 20 mg total protein was harvested for analysis after treatment with either DMSO or DRB for 30 minutes (first replicate: heavy cells DRB; light cells: DMSO; second replicate: heavy cells DMSO; light cells: DRB). After removing the media, each dish was scraped in 750 µl 95°C SDT (4% SDS, 100 mM Tris pH 7.9, 10 mM TCEP) buffer with subsequent heating at 95°C for 10 minutes. Lysates were sonicated for two minutes each. Protein concentrations were determined using a Bradford assay and samples were mixed 1:1 based on total protein concentrations. FASP was carried out in two 10 kDa MWCO filters with a 50 mM iodoacetamide alkylation step and proteins were digested in 2M urea with 2% wt/wt Lys- C (Wako) for 6 h and 2% modified trypsin (Promega) for 12 h at 37°C. FASP eluates were acidified and desalted on Oasis HLB extraction cartridges.

### TiO2 Phosphopeptide Enrichment, ERLIC Chromatography, and LC-MS/MS

Protocols were carried out as described (Stuart, Houel et al., 2015). An Orbitrap Velos (Thermo Fisher) was used for quantitative proteome analysis while an Orbitrap LTQ (Thermo Fisher) was used for phosphoproteomics. The samples were run on a 60 mins gradient / 10 HCD method.

### Proteomics data analysis

All raw mass spectrometry files for phosphoproteomics and quantitative proteomics were searched using the MaxQuant (v1.5.0.35) software package. Duplicate proteomic and phosphoproteomic were searched individually against the Uniprot human proteome database (downloaded on 16/01/2013) using the following MaxQuant parameters: multiplicity was set to 2 (heavy/light) with Arg10 and Lys8 selected, LysC/P was selected as an additional enzyme, “re-quantify” was unchecked, and Phospho (STY) was selected as a variable modification in both runs.

For phosphosite analysis, the Phospho (STY) table was processed with Perseus (v1.6.2.3) using the following workflow: reverse and contaminant reads were removed, the site table was expanded to accommodate differentially phosphorylated peptides, and rows without any quantification were removed after site table expansion. Normalized heavy to light ratios were log2 transformed for statistical analyses. Differential abundance of peptides following DRB treatment was estimated by t-tests with Welch correction, two sided, unpaired. The volcano plot was prepared with GraphPad Prism 9.1. Sequence motif plots were prepared with WebLogo 3 (Crooks, Hon et al., 2004).

### Generation of phosphoantibodies

Rabbit phosphoantibody against SF3B1 T142P was generated by Eurogentec based on the peptide sequence: H - CAD GGK T(PO3H2)PD PKM N - NH2 for SF3B1. Phosphoantibodies specificity was ensured by selecting against recognition of the unphosphorylated peptide.

## QUANTIFICATION AND STATISTICAL ANALYSIS

### Gene annotation

The Gencode V35 annotation, based on the hg38 version of the human genome, was used to extract the list of protein-coding genes. A list of 9,883 expressed protein-coding genes was obtained by keeping only the genes longer than 2 kb and with their highest transcript isoform expressed in two nuclear RNA-seq in HeLa cells (Nojima, Tellier et al., 2018) at more than 0.1 transcript per million (TPM), following quantification of transcript expression with Salmon version 0.14.1 (Patro, Duggal et al., 2017). The list of used exons was obtained by extracting the location of exons from Gencode V35 from the highest transcribed nuclear poly(A)+ RNA of each of the 9,883 protein-coding genes.

### mNET-seq data processing

Adapters were trimmed with Cutadapt version 1.18 (Martin, 2011) in paired-end mode with the following options: --minimum-length 10 -q 15,10 -j 16 – A GATCGTCGGACTGTAGAACTCTGAAC – a AGATCGGAAGAGCACACGTCTGAACTCCAGTCAC. Trimmed reads were mapped to the human GRCh38.p13 reference sequence with STAR version 2.7.3a (Dobin, Davis et al., 2013) and the parameters: --runThreadN 16 -- readFilesCommand gunzip -c -k --limitBAMsortRAM 20000000000 --outSAMtype BAM SortedByCoordinate. SAMtools version 1.9 (Li, Handsaker et al., 2009) was used to retain the properly paired and mapped reads (-f 3). A custom python script (Nojima et al., 2015) was used to obtain the 3′ nucleotide of the second read and the strandedness of the first read. Strand-specific bam files were generated with SAMtools. Samples normalization was checked against the termination region of the *RNU2* snRNA, which is known to be insensitive to DRB (Medlin, Uguen et al., 2003), and also verified for consistency against a previously published GRO-seq performed by our group in HeLa cells treated for 30 minutes with DMSO or DRB (Laitem et al., 2015). FPKM-normalized bigwig files were created with deepTools version 3.4.2 (Ramirez, Ryan et al., 2016) bamCoverage tool with the parameters -bs 1 -p max –normalizeUsing RPKM.

### Spike-in ChIP-seq processing

Adapters were trimmed with Cutadapt with the following options: --minimum-length 10 -q 15, 10 -j 16 -a AGATCGGAAGAGCACACGTCTGAACTCCAGTCA. Trimmed reads were mapped to the human GRCh38.p13 and to the mouse GRCm38 reference genomes with STAR and the parameters: --runThreadN 16 --readFilesCommand gunzip -c -k –alignIntronMax 1 -- limitBAMsortRAM 20000000000 --outSAMtype BAM SortedByCoordinate. SAMtools was used to retain the properly mapped reads and to remove PCR duplicates. Reads mapping to the DAC Exclusion List Regions (accession: ENCSR636HFF) were removed with BEDtools version 2.29.2 (Quinlan & Hall, 2010). SAMtools view with the –s option was used to subsample all the bam files to the bam file containing the lowest number of reads. The normalization factor was then calculated as: (number of mouse reads) / (number of mouse + number of human reads)) and applied to the generation of the bigwig files with deepTools bamCoverage tool with the parameters -bs 10 -p max –e –scaleFactor.

### Analysis of 3’READS data

The analysis was performed exactly as in (Neve et al., 2016). Briefly, raw reads were mapped to the human genome (hg19) with Bowtie 2 (Langmead & Salzberg, 2012) using the option “- M 8 --local”. Reads that were shorter than 15 nt, were non-uniquely mapped to genome (MAPQ < 10), or contained more than 2 mismatches in alignment were discarded. FPKM-normalized bigwig files were created with deepTools bamCoverage tool with the parameters -bs 10 -p max --normalizeUsing RPKM.

### Metagene profiles

Metagene profiles of genes scaled to the same length were then generated with Deeptools2 computeMatrix tool with a bin size of 10 bp and the plotting data obtained with plotProfile – outFileNameData tool. Graphs representing the (IP – Input) signal (ChIP-seq) or the mNET-seq signal were then created with GraphPad Prism 9.1. Metagene profiles are shown as the average of two biological replicates.

### P-values and significance tests

P-values were computed with an unpaired two-tailed Student’s t test. Statistical tests were performed in GraphPad Prism 9.1.

## Acknowledgments

We thank Chris Norbury for helpful comments on the manuscript. We thank Shabaz Mohammed for help with the SILAC phosphoproteomics. We thank the High-Throughput Genomics Group at the Wellcome Trust Centre for Human Genetics for sequencing. This work was supported by a Wellcome Trust Investigator Award (WT106134AIA and WT210641/Z/18/Z) and a Biotechnology and Biological Sciences Research Council grant (BB/R016836/1) to S.M. A.F.’s research is funded by the BBSRC (BB/N001184/1) and J.N. is funded by an MRC studentship.

## Competing interests

The authors declare that they have no conflict of interest.

## Data availability

Sequencing data have been deposited in GEO under accession code GSE176541. The mass spectrometry proteomics data have been deposited to the ProteomeXchange Consortium via the PRIDE (Perez-Riverol, Csordas et al., 2019) partner repository with the dataset identifier PXD026720. All data generated or analysed during this study are included in the manuscript and supporting files. We include full excel spreadsheets representing original mass spectrometry data.

## Author contributions

**Michael Tellier:** Conceptualization; Data curation; Formal analysis; Investigation; Validation; Visualization; Writing—original draft; Writing—review & editing. **Justyna Zaborowska:** Investigation. **Jonathan Neve:** Investigation. **Takayuki Nojima:** Investigation. **Svenja Hester:** Investigation. **Marjorie Fournier:** Investigation; Formal analysis. **Andre Furger:** Funding acquisition; Supervision. **Shona Murphy:** Conceptualization; Funding acquisition; Investigation; Project administration; Supervision; Writing—original draft; Writing—review & editing.

In addition to the CRediT author contributions listed above, the contributions in detail are: MT performed all the experiments and bioinformatics analysis except for the following: initial help with the mNET-seq from JZ and TN, JN performed the 3’READS libraries preparation and sequencing, SH processed the phosphoproteomics samples, and MF processed and analysed the SF3B1 proteomics samples. AF supervised JN. SM made the CDK9as cell line, carried out the cell proliferation analyses and supervised MT and JZ. MT and SM wrote the manuscript.

---

**Figure EV1.**
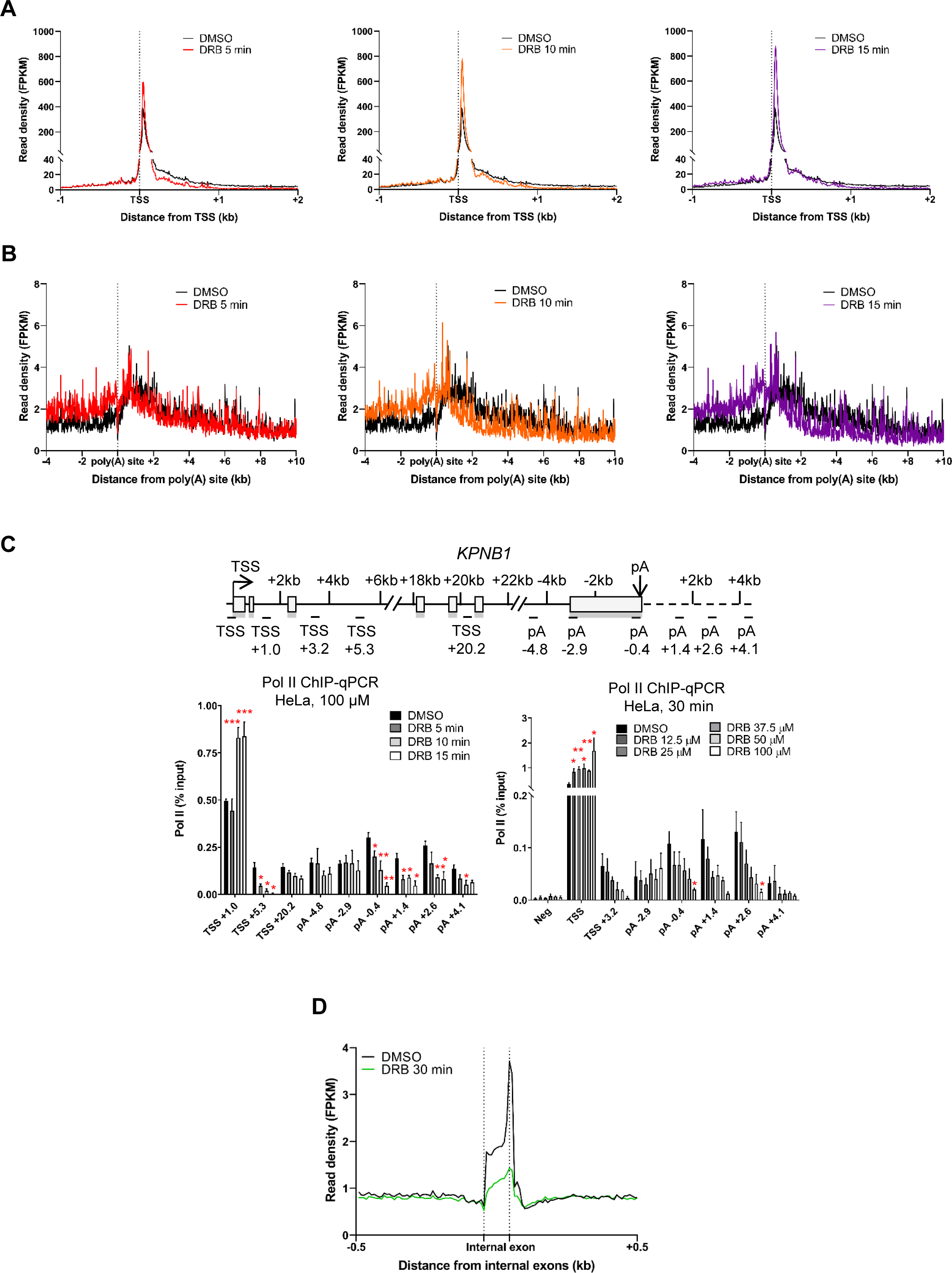
CDK9 inhibition causes an elongation defect starting at the last exon of protein-coding genes. **A.** Metagene profile of total pol II after treatment with DMSO (black) or 5 (red), 10 (orange), or 15 (purple) minutes with DRB (green) around the TSS of expressed protein-coding genes (n=6,965). **B.** Metagene profile of total pol II after treatment with DMSO (black) or 5 (red), 10 (orange), or 15 (purple) minutes with DRB (green) around the TSS of expressed protein-coding genes longer than 40 kb (n=2,816). **C.** ChIP-qPCR of total pol II with different treatment times or different concentrations of DRB or SNS-032 on KPNB1. n=3 biological replicates, mean ± SEM, p-value: * p < 0.05, ** p < 0.01, *** p < 0.001. Statistical test: two-tailed unpaired t test. **D.** Metagene profile of total pol II after 30 minutes treatment with DMSO (black) or DRB (green) around internal exons, not including first, penultimate or last exons, of expressed protein-coding genes (n=26,094).

**Figure EV2.**
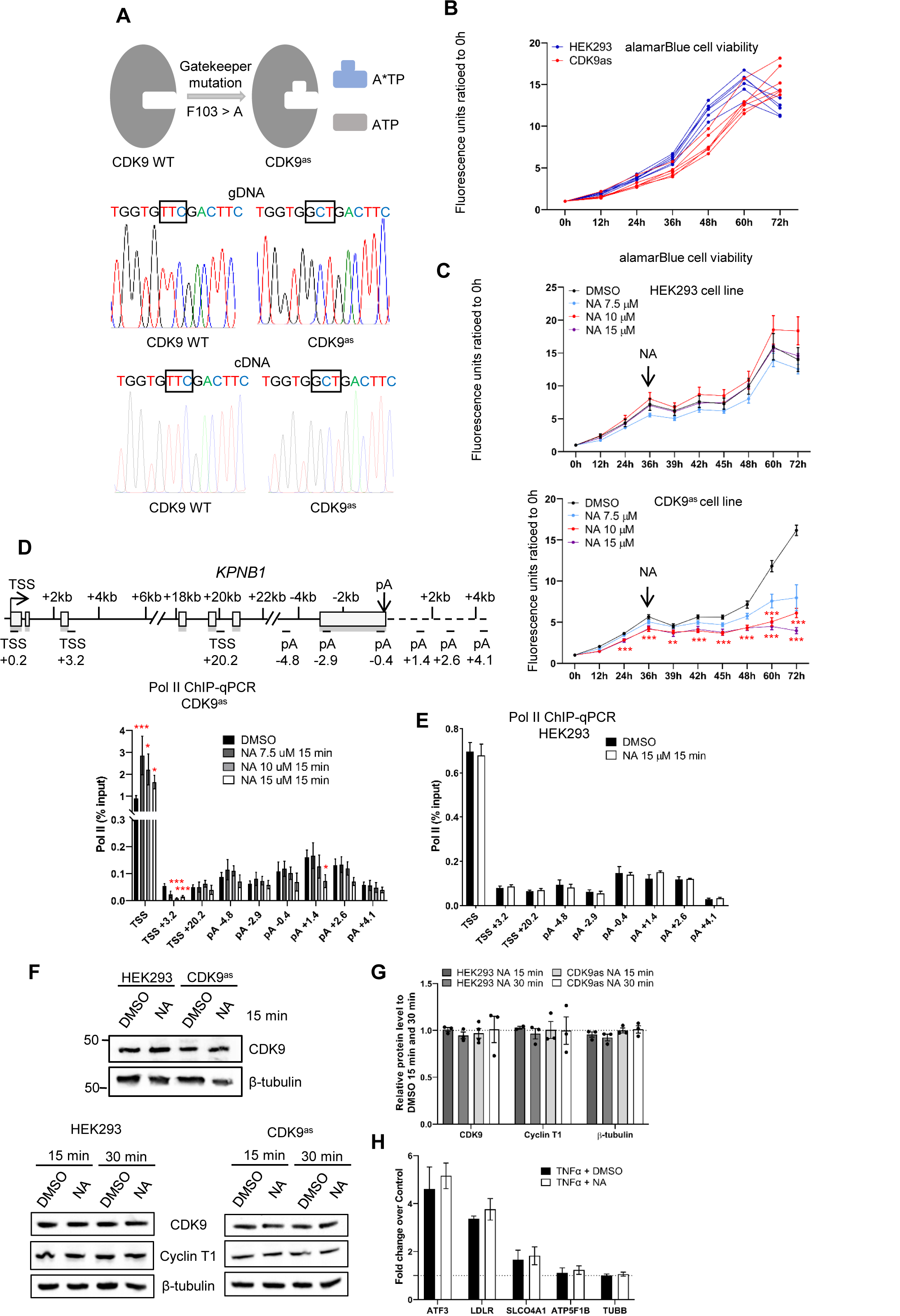
Inhibition of analog-sensitive (as) CDK9 produces similar results to small molecule CDK9 inhibitors. **A.** Schematic of the genome editing of the CDK9as cell line. **B.** alamarBlue cell viability of wild-type HEK293 and CDK9as cells. Each line represents a biological replicate. **C.** alamarBlueHS cell viability assay of wild-type HEK293 and CDK9as cells with different concentrations of 1-NA-PP1 added at the 36-hour time point. n=3 biological replicates, p-value: ** p < 0.01, *** p < 0.001. Statistical test: paired t-test with FDR multiple testing correction. **D.** ChIP-qPCR of total pol II with different concentrations of 1-NA-PP1 in CDK9as cells on KPNB1. n=3 biological replicates, mean ± SEM, p-value: * p < 0.05, *** p < 0.001. Statistical test: two-tailed unpaired t test. **E.** ChIP-qPCR of total pol II treated with 1-NA-PP1 in wild-type HEK293 cells on KPNB1. n=3 biological replicates, mean ± SEM, p-value: n.s. not significant. Statistical test: two-tailed unpaired t test. **F.** Western blot of CDK9, Cyclin T1, and β- tubulin as a loading control, on whole cell extracts of wild-type HEK293 and the CDK9as cell line treated with DMSO or 1-NA-PP1 for 15 or 30 minutes. **G.** Quantification of the western blots shown in F. n=2 biological replicates, mean ± SEM, p-value: not significant. Statistical test: two-tailed unpaired t test. **H.** qRT-PCR of nuclear polyadenylated mRNAs of several TNFα-induced or non-induced genes with a 30 minutes DMSO or a NA treatment. n=3 biological replicates, mean ± SEM, p-value: not significant. Statistical test: two-tailed unpaired t test.

**Figure EV3.**
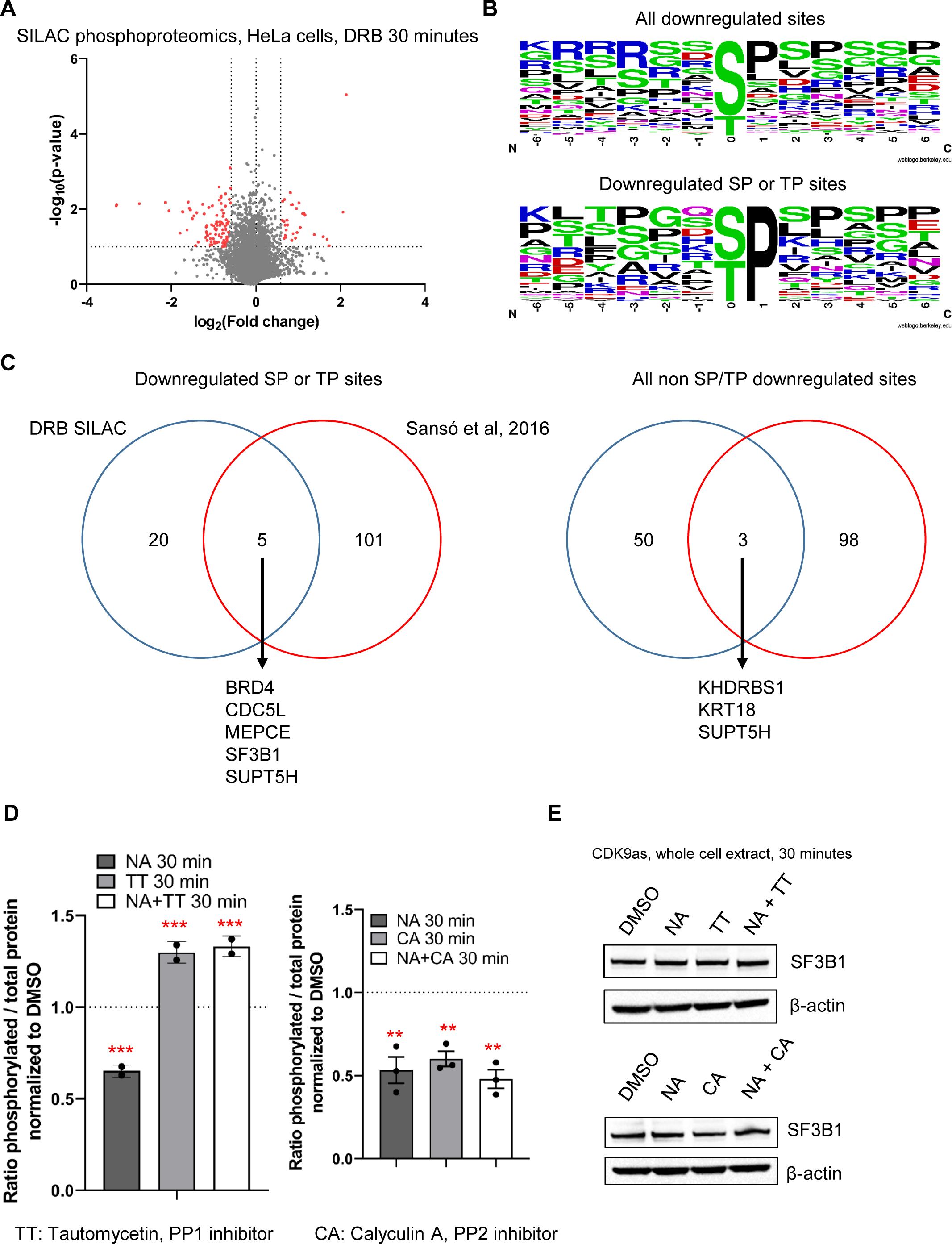
CDK9 phosphorylates several transcription and splicing factors in vivo. **A.** Volcano plot of SILAC phosphoproteomics in HeLa cells treated or not with 100 µM DRB for 30 minutes (in red: fold change > 1.5 in both biological duplicates, p-value < 0.1). **B.** Motif found around all the phosphorylation sites decreased following CDK9 inhibition of only the phosphorylation sites containing a ST or TP sites. **C.** Overlap between the proteins found to have at least one phosphopeptides decreased in our study versus an alternative experimental strategy used to identify CDK9 targets in cell extracts (Sanso et al., 2016). **D.** Quantification of the western blots shown in Figure 4B. n=2 biological replicates for the TT set, n=3 biological replicates for the CA set, mean ± SEM, p-value: ** p < 0.01, *** p < 0.001. Statistical test: two-tailed unpaired t test. **E.** Western blot of SF3B1 and β-actin, as a loading control, on the whole cell extract of CDK9as cells after 30 minutes DMSO, NA, CA, or NA+CA treatment, or DMSO, NA, TT, NA+TT treatment.

**Figure EV4.**
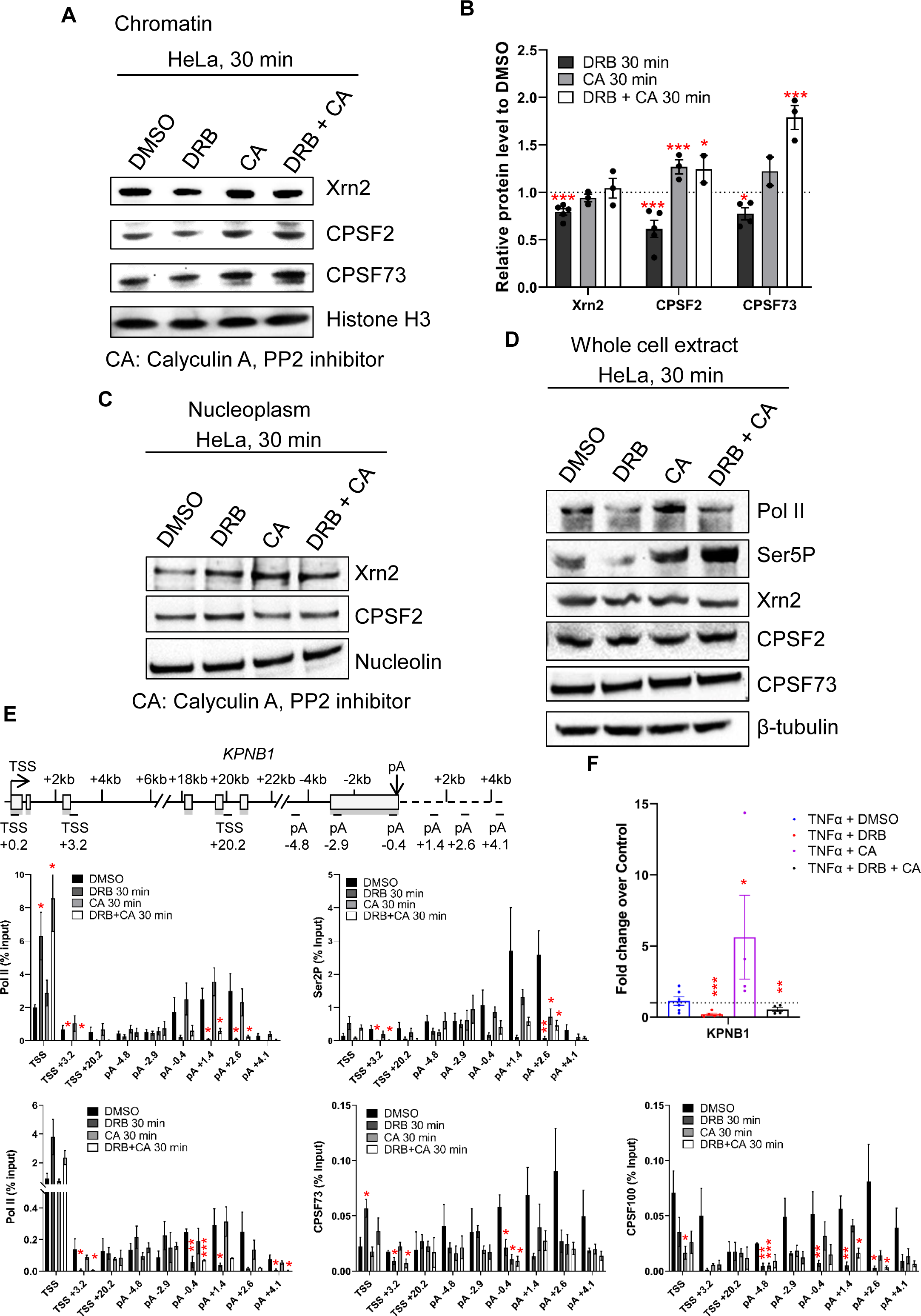
CDK9 and PP2A regulate mRNA cleavage and polyadenylation. **A.** Western blot of Xrn2, CPSF2, CPSF73, and histone H3 as a loading control, on the chromatin fraction of HeLa cells after 30 minutes DMSO, DRB, CA, or DRB+CA treatment. **B.** Quantification of the western blots shown in C. n=3 biological replicates, mean ± SEM, p-value: * p < 0.05, ** p < 0.01, *** p < 0.001. Statistical test: two-tailed unpaired t test. **C.** Western blot of Xrn2, CPSF2, and Nucleolin as a loading control, on the nucleoplasm fraction of HeLa cells after a 30 minutes DMSO, DRB, CA, or DRB+CA treatment. The CPSF73 antibody is not shown at it does not provide reliable results on the nucleoplasm fraction. **D.** Western blot of total pol II, Ser5P, Xrn2, CPSF2, CPSF73, and β-tubulin as a loading control, on whole cell extract of HeLa cells treated for 30 minutes with DMSO, DRB, CA, or DRB+CA. **E.** ChIP-qPCR of pol II, Ser2P, CPSF73, or CPSF2 after 30 minutes treatment with DMSO, DRB, CA, or DRB+CA on KPNB1. n=3 biological replicates, mean ± SEM, p- value: * p < 0.05, ** p < 0.01, *** p < 0.001. Statistical test: two-tailed unpaired t test. **F.** qRT-PCR of nuclear polyadenylated mRNAs of the KPNB1 gene with a 30 minutes DMSO, DRB, TT, CA, DRB+TT, or DRB+CA treatment. n=4 biological replicates, mean ± SEM, p-value: * p < 0.05, ** p < 0.01, *** p < 0.001. Statistical test: two-tailed unpaired t test.

**Figure EV5.**
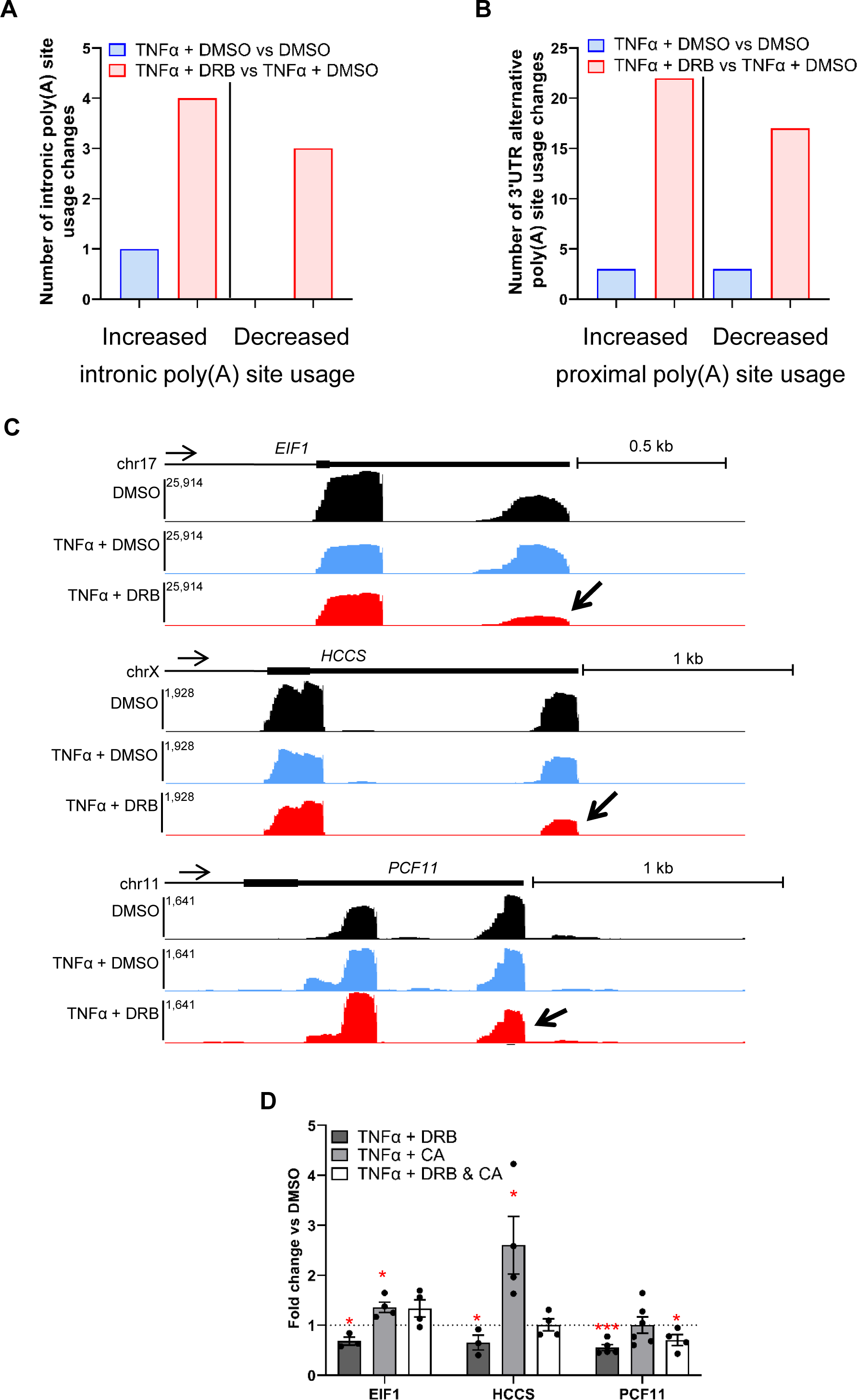
CDK9 and PP2A are involved in mRNA alternative poly(A) site usage. **A.** Number of genes undergoing significant increased or decreased intronic poly(A) site usage in both biological replicates of the 3’READS experiments. **B.** Number of genes undergoing significant increased or decreased proximal poly(A) site usage in both biological replicates of the 3’READS experiments. **C.** Screenshots of the genome browser 3’READS tracks at the 3’end of protein-coding genes EIF1, HCCS, and PCF11, which are undergoing decreased poly(A) site usage following CDK9 inhibition with DRB (indicated by the arrow). **D.** qRT-PCR of nuclear polyadenylated mRNAs of the EIF1, HCCS, and PCF11 genes gene with a 30 minutes DMSO, DRB, CA, or DRB+CA treatment. Two pairs of primers were used for each gene, one pair for the total transcripts level and one pair specific for the transcript using the distal poly(A) site usage. The data are shown as distal poly(A) site / total and normalized to the DMSO control. A value below 1 corresponds to a shift to proximal poly(A) site usage while a value superior to 1 corresponds to an increased distal poly(A) site usage. n=3 biological replicates, mean ± SEM, p-value: * p < 0.05, *** p < 0.001. Statistical test: two-tailed unpaired t test.

## Notes

### Competing Interest Statement

The authors have declared no competing interest.

