## Appendix Figures and Supplementary Tables 1 and 2 for "CDK9 and PP2A regulate RNA polymerase II transcription termination and coupled RNA maturation"

### Table of contents

**Appendix Figure S1.** CDK9 inhibition abrogates polyadenylation……………………………………2

**Appendix Figure S2.** Inhibition of a CDK9 analog sensitive (as) cell line produces similar results to the current CDK9 inhibitors……………………………………………………………………...4

**Appendix Figure S3.** CDK9 phosphorylates several transcription and splicing factors *in vivo*……6

**Appendix Figure S4.** PP1 and PP2A regulates transcription of protein-coding genes……………...8

**Appendix Figure S5.** CDK9 and PP2A regulate mRNA cleavage and polyadenylation…………...10

**Appendix Table 1:** List of antibodies used in mNET-seq, ChIP, co-immunoprecipitation, and western blots……………………………………………………………………………………..12

**Appendix Table 2:** List of primers used for ChIP-qPCR and qRT-PCR……………………………..13

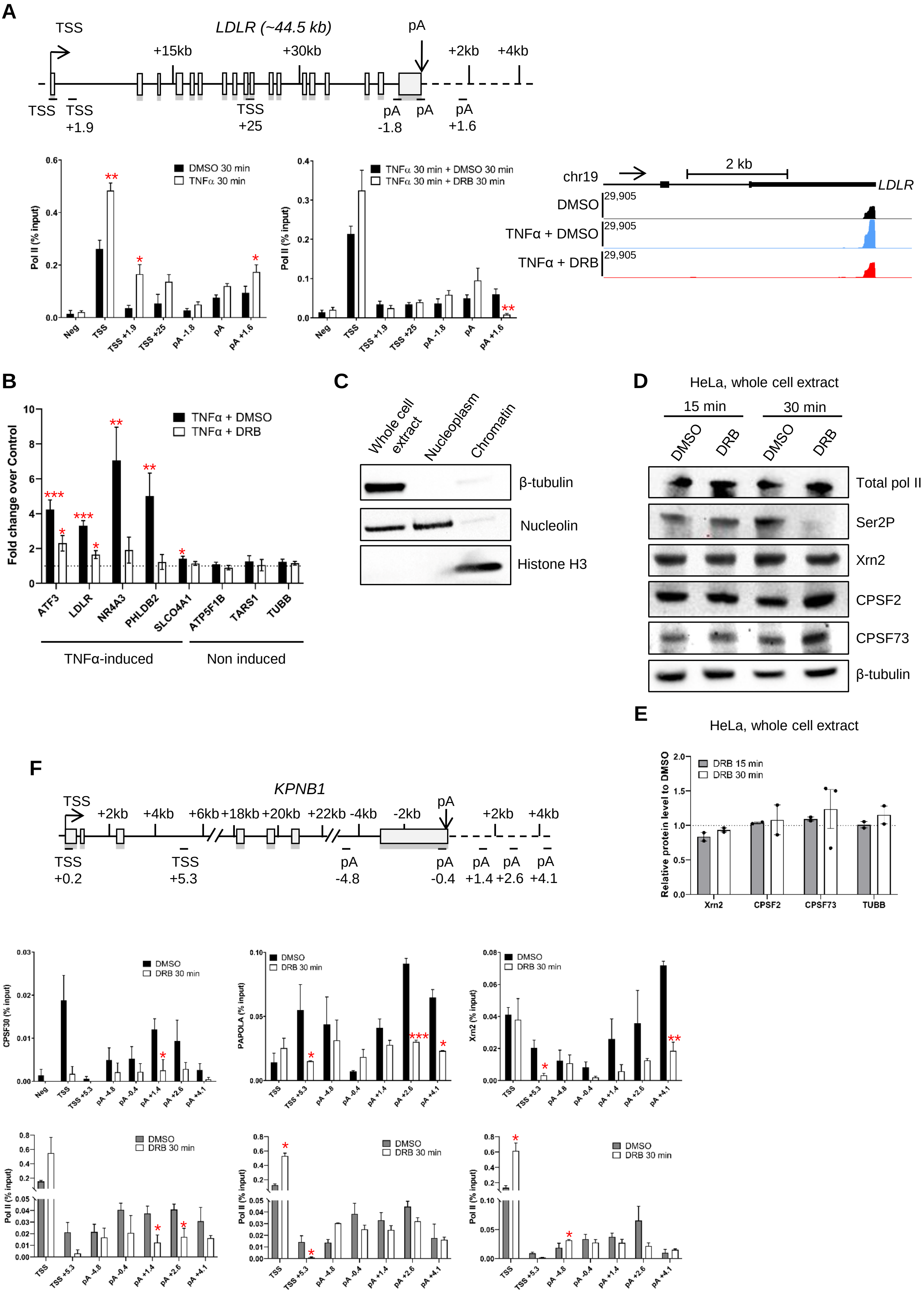


Appendix Figure S1

**Appendix Figure S1. CDK9 inhibition abrogates polyadenylation.**

**A.** Pol II ChIP-qPCR on the TNFα-induced gene LDLR and screenshot of the genome browser track of the 3'READS experiments for this gene. n=3 biological replicates, mean ± SEM, p-value: * $p < 0.05$, ** $p < 0.01$. Statistical test: two-tailed unpaired t test. **B.** qRT-PCR of nuclear polyadenylated mRNAs of several TNFα-induced or non-induced genes with a 30 minutes DMSO or a DRB (100 µM here and in all other figures unless stated otherwise) treatment. n=5 biological replicates, mean ± SEM, p-value: * $p < 0.05$, ** $p < 0.01$, *** $p < 0.001$. Statistical test: two-tailed unpaired t test. **C.** Western blot of β-tubulin, Nucleolin, and Histone H3 on whole cell extract, nucleoplasm, and chromatin fractions. **D**. Western blot of total pol II, Ser2P, Xrn2, CPSF2, CPSF73, and β-tubulin as a loading control, on whole cell extract. **E.** Quantification of the western blots shown in D. n=2 biological replicates, mean ± SEM, p-value: n.s. not significant. Statistical test: two-tailed unpaired t test **F.** ChIP-qPCR of total pol II, CPSF73, PAPOLA, and Xrn2 on the model gene, *KPN.B1*. n=3 biological replicates, mean ± SEM, p-value: * $p < 0.05$, ** $p < 0.01$, *** $p < 0.001$. Statistical test: two-tailed unpaired t test.

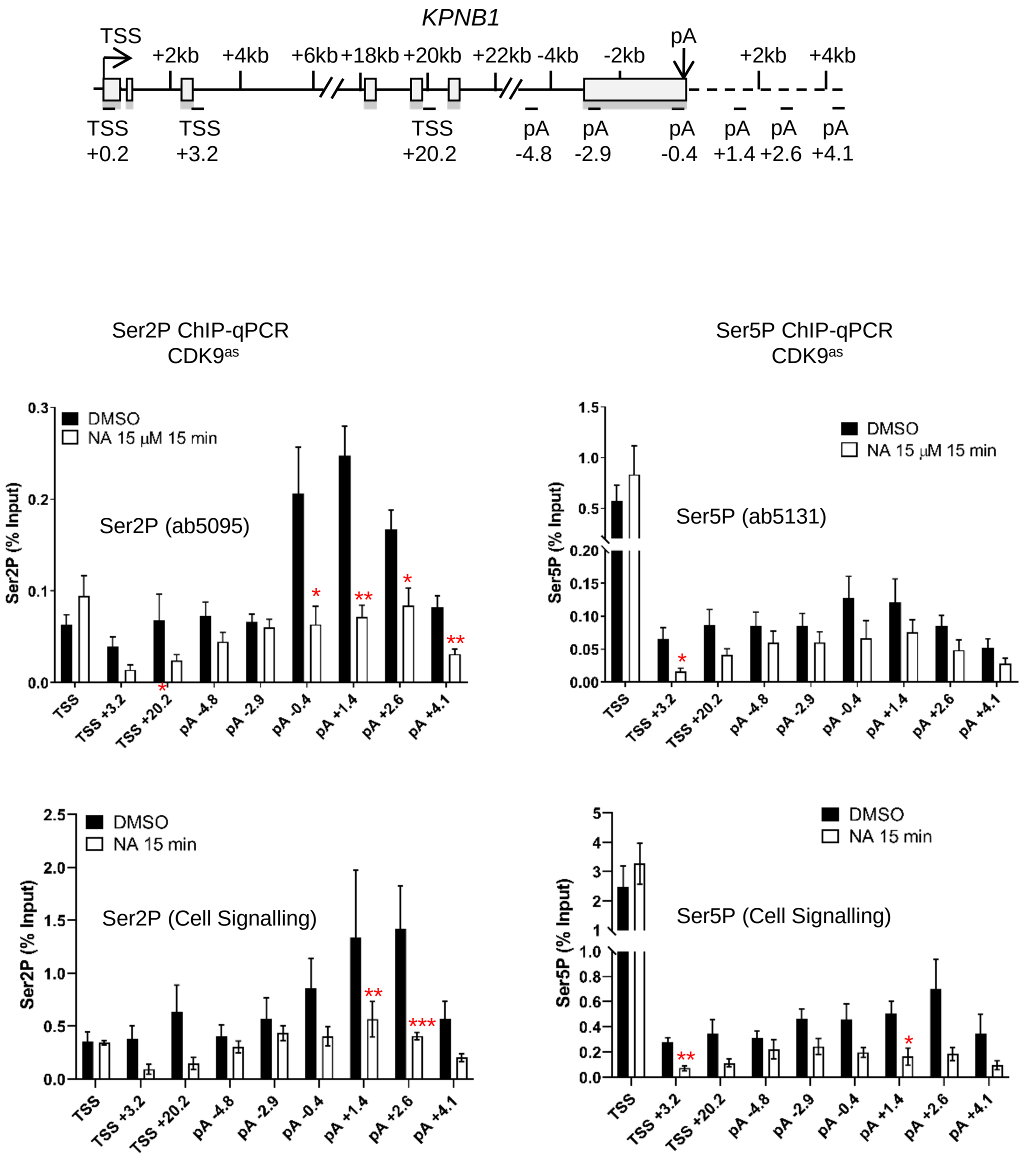


Appendix Figure S2

**Appendix Figure S2. Inhibition of a CDK9 analog sensitive (as) cell line produces similar results to the current CDK9 inhibitors.**
ChIP-qPCR of Ser2P (Abcam or Cell Signaling), and Ser5P (Abcam or Cell Signaling) in CDK9as cells treated with 15mM 1-NA-PP1 for 15 minutes on *KPNB1*. n=3 biological replicates, mean ± SEM, p-value: * $p < 0.05$, ** $p < 0.01$, *** $p < 0.001$. Statistical test: two-tailed unpaired t test.

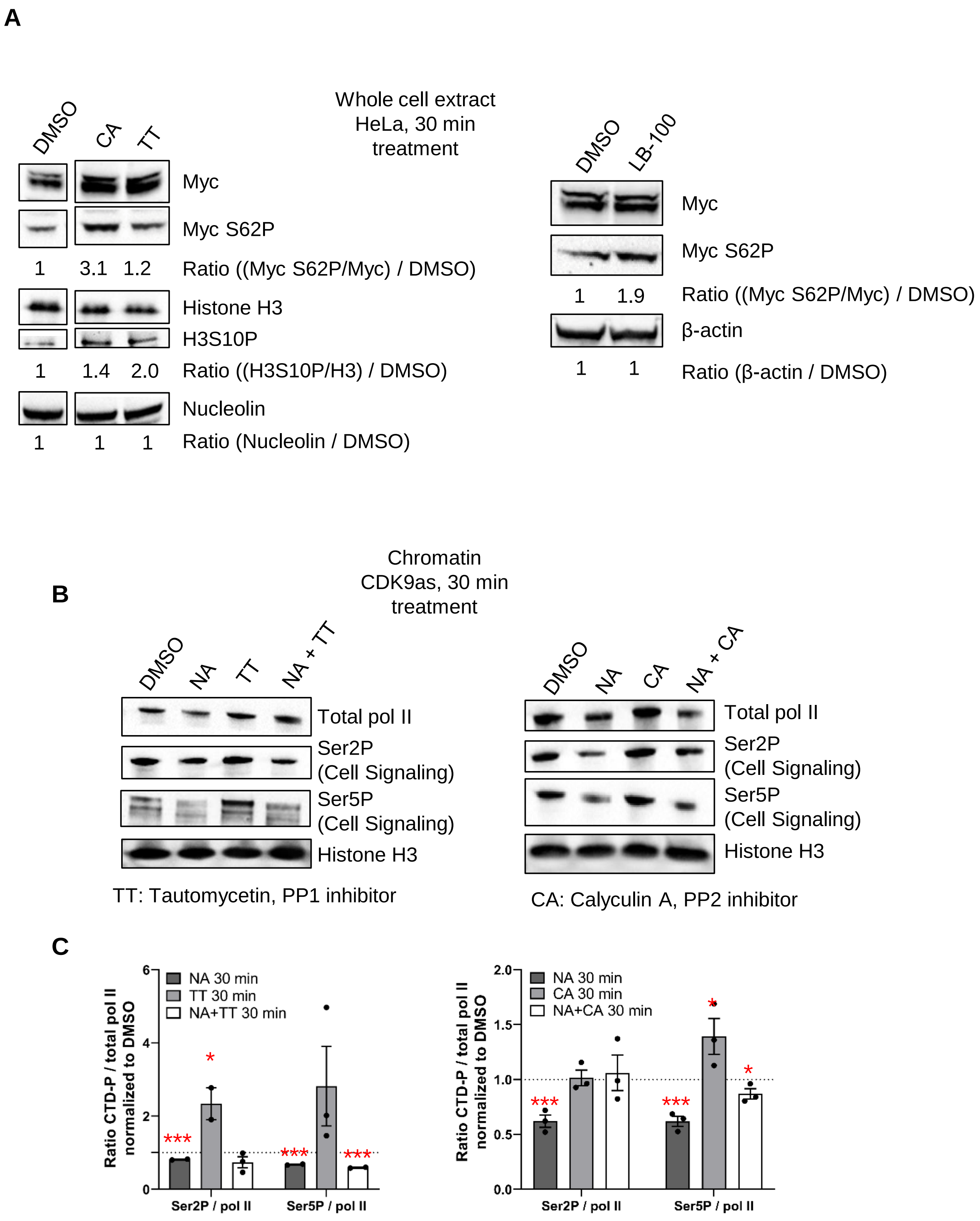


Appendix Figure S3

**Appendix Figure S3. CDK9 phosphorylates several transcription and splicing factors in vivo.**

**A.** Western blot of Myc, Myc S62P, Histone H3, Histone H3 S10P, and Nucleolin as a loading control, on whole cell extract of HeLa cells treated for 30 minutes with DMSO, DRB, CA, LB-100, or TT. **B.** Western blot of total pol II, Ser2P, Ser5P, and histone H3 as a loading control, on the chromatin fraction of CDK9as cells treated for 30 minutes with DMSO, NA, CA, TT, NA+CA, or NA+TT. **C.** Quantification of the western blots shown in B. n=2 biological replicates, mean ± SEM, p-value: * $p < 0.05$, *** $p < 0.001$. Statistical test: two-tailed unpaired t test.

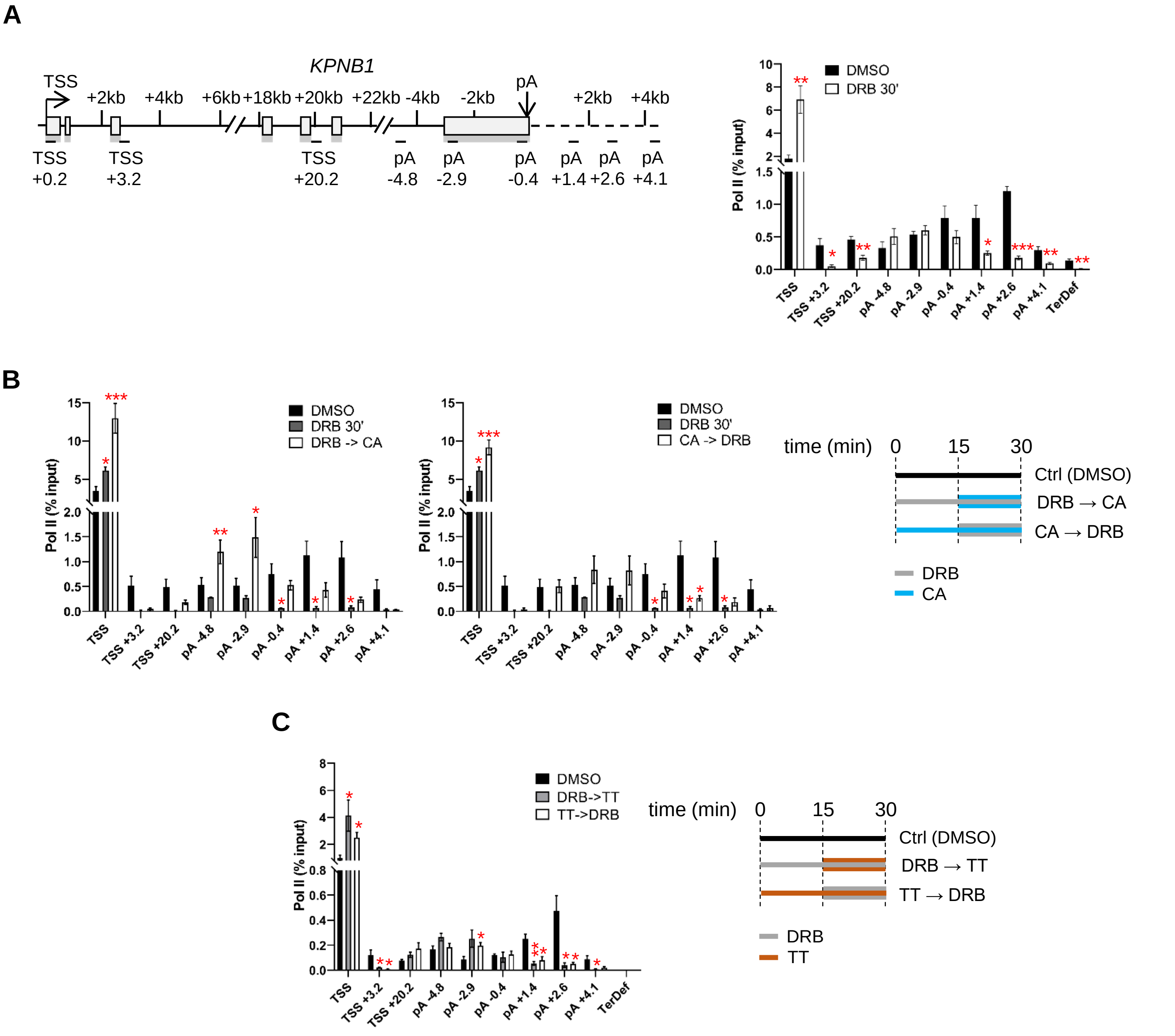


Appendix Figure S4

**Appendix Figure S4. PP1 and PP2A regulates transcription of protein-coding genes.**

**A.** ChIP-qPCR of total pol II after 30 minutes treatment with DMSO or DRB on *KPNB1*. n=3 biological replicates, mean ± SEM, p-value: * $p < 0.05$, ** $p < 0.01$, *** $p < 0.001$. Statistical test: two-tailed unpaired t test. **B.** ChIP-qPCR of total pol II after 30 minutes treatment with DMSO or DRB, or 15 minutes treatment with DRB or CA followed by another 15 minutes with CA (DRB->CA) or DRB (CA->DRB), respectively, on *KPNB1*. n=3 biological replicates, mean ± SEM, p-value: * $p < 0.05$, ** $p < 0.01$, *** $p < 0.001$. Statistical test: two-tailed unpaired t test. **C.** ChIP-qPCR of total pol II after 30 minutes treatment with DMSO or DRB, or 15 minutes treatment with DRB or TT followed by another 15 minutes with TT (DRB->TT) or DRB (TT->DRB), respectively, on *KPNB1*. n=3 biological replicates, mean ± SEM, p-value: * $p < 0.05$, ** $p < 0.01$, *** $p < 0.001$. Statistical test: two-tailed unpaired t test.

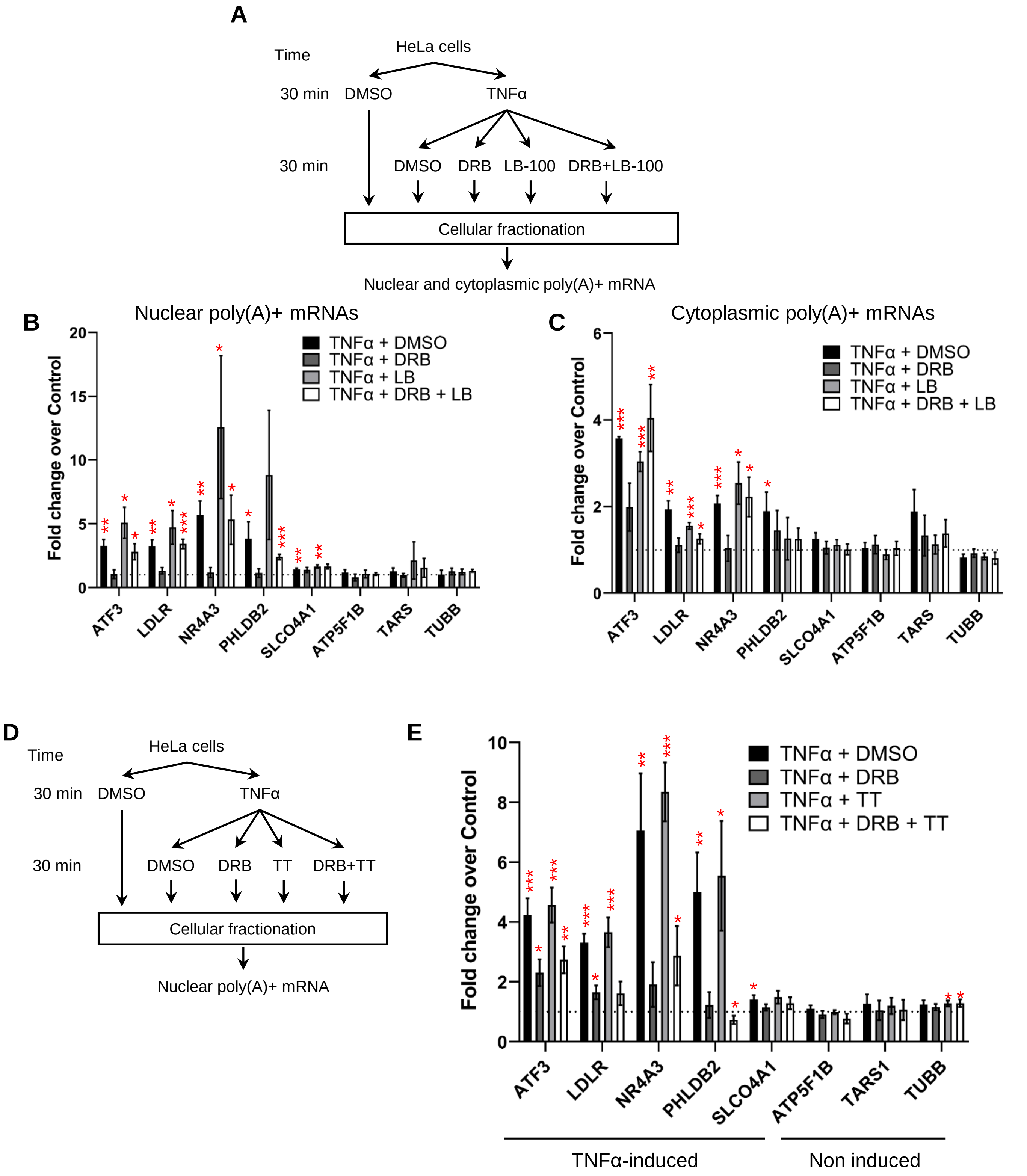


Appendix Figure S5

**Appendix Figure S5. CDK9 and PP2A regulate mRNA cleavage and polyadenylation.**

**A.** Schematic of the nuclear and cytoplasmic qRT-PCR experiments. **B.** qRT-PCR of nuclear polyadenylated mRNAs of several TNFα induced or non-induced genes with a 30 minutes DMSO, DRB, LB, or DRB+LB-100 treatment. n=3 biological replicates, mean ± SEM, p-value: * $p < 0.05$, ** $p < 0.01$, *** $p < 0.001$. Statistical test: two-tailed unpaired t test. **C.** qRT-PCR of cytoplasmic polyadenylated mRNAs of several TNFα induced or non-induced genes with a 30 minutes DMSO, DRB, LB-100, or DRB+LB-100 treatment. n=3 biological replicates, mean ± SEM, p-value: * $p < 0.05$, ** $p < 0.01$, *** $p < 0.001$. Statistical test: two-tailed unpaired t test. **D.** Schematic of the nuclear qRT-PCR experiments. **E.** qRT-PCR of nuclear polyadenylated mRNAs of several TNFα induced or non-induced genes with a 30 minutes DMSO, DRB, TT, or DRB+TT treatment. n=4 biological replicates, mean ± SEM, p-value: * $p < 0.05$, ** $p < 0.01$, *** $p < 0.001$. Statistical test: two-tailed unpaired t test.

**Appendix Table 1:** List of antibodies used in mNET-seq, ChIP, co-immunoprecipitation, and western blots.

| Antibody | Supplier | Catalogue number | Applications |
|---|---|---|---|
| Mouse monoclonal anti-Pol II CTD, Total | MBL international | MABI0601 | mNET-seq |
| Rpb1 NTD (D8L4Y) Rabbit mAb | Cell Signaling Technology | 14958S | ChIP, ChIP-seq, Western blot |
| Phospho-Rpb1 CTD (Ser2) (E1Z3G) Rabbit mAb | Cell Signaling Technology | 13499S | ChIP, Western blot |
| Phospho-Rpb1 CTD (Ser5) (D9N5I) Rabbit mAb | Cell Signaling Technology | 13523S | ChIP, Western blot |
| RNA pol II antibody (mAb) | Active Motif | 39097 | Co-immunoprecipitation |
| Anti-RNA polymerase II CTD repeat YSPTSPS (phospho S2) antibody | Abcam | ab5095 | Western blot |
| Anti-RNA polymerase II CTD repeat YSPTSPS (phospho S5) antibody | Abcam | ab5131 | Western blot |
| Rabbit anti-SF3b155/SAP155 Antibody, Affinity Purified | Bethyl Laboratories | A300-996A | Western blot |
| SF3b1 T142P | This study | / | Western blot |
| Anti-SAP155 Monoclonal Antibody | MBL international | D221-3 | Co-immunoprecipitation |
| Rabbit anti-SF3B3 Antibody, Affinity Purified | Bethyl Laboratories | A302-508A | Western blot |
| Myc | Santa Cruz Biotechnology | sc-764X | Western blot |
| Myc S62P | Abcam | ab51156 | Western blot |
| Anti-Cdk9 antibody [EPR3119Y] | Abcam | ab76320 | Western blot |
| Cyclin T1 (H-245) | Santa Cruz Biotechnology | sc-10750 | Western blot |
| Rabbit anti-CPSF30 Antibody, Affinity Purified | Bethyl Laboratories | A301-584A | ChIP, Western blot |
| Rabbit anti-CPSF73 Antibody, Affinity Purified | Bethyl Laboratories | A301-091A | ChIP, Western blot |
| Rabbit anti-CPSF100 Antibody, Affinity Purified | Bethyl Laboratories | A301-581A | ChIP, Co-immunoprecipitation, Western blot |
| Rabbit anti-PAPOLA Antibody, Affinity Purified | Bethyl Laboratories | A301-008A | ChIP, Western blot |
| Rabbit anti-XRN2 Antibody, Affinity Purified | Bethyl Laboratories | A301-103A | ChIP, Western blot |
| Histone H3 | Abcam | ab1791 | Western blot |
| Histone H3 S10P | Life Technologies | PA517869 | Western blot |
| Nucleolin | Abcam | ab22758 | Western blot |
| Beta-actin | Cell Signaling Technology | 4967S | Western blot |
| Anti-beta Tubulin antibody - Loading Control | Abcam | ab6046 | Western blot |
| Normal Rabbit IgG | Cell Signaling Technology | 2729S | ChIP, Co-immunoprecipitation |

**Appendix Table 2:** List of primers used for ChIP-qPCR and qRT-PCR

| Name | Forward primer | Reverse primer |
|---|---|---|
| **ChIP primers** | | |
| KPNB1 TSS | TTACTTCCTCCCTCCAAATGGG | ACAGCCTCCCTTCCTTCTTTC |
| KPNB1 TSS+3.2 | GCCCAGAGAACAAGAAATCG | GGAATGGACAAGCTGTGTTG |
| KPNB1 TSS+20.2 | TGCAAGAGCCAGTGGGAACACTT | CCTCTACTCAGCAATGATACTTC |
| KPNB1 pA-4.8 | CTGAGGAAACTGAAGAACCAAG | GAAGGCAGTGCTTGCCAGAAT |
| KPNB1 pA-2.9 | GAGGAGTGTGCACGGATGCTGAA | CCAAGATGGCCGATGTTATGG |
| KPNB1 pA-0.4 | TAGTTACCGTCTGCTTGGGAAGATG | CCTCTGACAGCAAGTCCAACATT |
| KPNB1 pA+1.4 | GACTCATCACACCAAGGTCAC | GATAGTGCTGGGAAGGAAATGG |
| KPNB1 pA+2.6 | GTACATCTCAGCTTTGGCATATG | GCCCAGAACATAGCAGGCATTGC |
| KPNB1 pA+4.1 | GTTTCACCGTGTTAGCCAGGATGG | CCACAGCCATGTTCATTTCTGC |
| KPNB1 TD | AGGAGCATGGCTTTTCTCTG | TCATGCTGGAACTGGTTGAG |
| Neg | TGGTACAACCACAGCTCAGTG | AAGCTGGACATGGTTGTGTG |
| LDLR TSS | AATCACCCCACTGCAAACTC | TAGCTGGAAACCCTGGCTTC |
| LDLR TSS +1.9 | TGGGATTGCCTGATGAACAC | AAGGCAGTTCAAAGCTCTCC |
| LDLR TSS +25 | TTCAGTGTGGTGCTGACAAC | TTCTCTGCTGGAAACCCAAC |
| LDLR pA -1.8 | CAGAGAAGACCAAAGCATTGCC | AATCCCAACCCAAGCCATTG |
| LDLR pA | AATCGCCGTGTTACTGTTGC | TGCCAATCCCTTGTGACATC |
| LDLR pA +1.6 | GTGATTGTGTTCTCTGCTGTCG | TCGCACTTAGCACTCAACAG |
| **qRT-PCR primers** | | |
| ATF3 | AACCACAGTCAGTGGAGAGATG | TTCTCACAGCTGCAAACACC |
| LDLR | TTTGACGGGACTTCAGGTTC | TCCCTTGTGACATCTTCACG |
| NR4A3 | AAGCCACCAGCTGTTAATGG | GCAATGCTGTTAGAGGAGCAG |
| PHLDB2 | GAAACGACTTCAGGCAAGTCTC | GCCATGTTTTAGGAAAGAGCAC |
| SLCO4A1 | TCCTCTTCTTTGCCATAGCC | ACAAGTTTCCAGGCCATCTG |
| ATP5F1B | ACCCATTGAAGAAGCTGTGG | CAATCAAGGCTCTTGTGCAG |
| TARS1 | CATGGAAAAGGAGGAACAGC | CTTTGCCAAACTCCTCCAAG |
| TUBB | ATATGTTCCTCGTGCCATCC | TTTGGCCCAGTTGTTACCTG |
| GAPDH | CAACGACCACTTTGTCAAGC | TTCCTCTTGTGCTCTTGCTG |
| EIF1 distal pA | GCCTGAAACCAAGCAATACC | GTTCCGGCCATAGTTGTTTG |
| HCCS distal pA | CTCGAAAAGCCTGAACAACC | TCTTGAGCATCACCATGGAC |
| PCF11 distal pA | TGGAATTGAACAGCAACCTC | GAACCCCCTTTTGAAGGATG |
| EIF1 total | AGGGATCGCTGATGATTACG | TCTCCATATTCCGGATGCTC |
| HCCS total | TGGTGGTGAAGTCAACAAGG | CCACCAAGCGACTTTCATTC |
| PCF11 total | TCCACTCCTCCAATTGTTCC | AGCTCCAGCTTTTTCTGCTG |
